# Fatty acid desaturation guides cellular decisions between ferroptosis and cellular senescence

**DOI:** 10.1101/2024.10.03.616257

**Authors:** Chisaka Kuehnemann, Samantha Jezak, Bronwyn Mogck, Niharika Kura, Yuheng Che, Maija Vaittinen, Ioannis Siokas, Rushmeen Tariq, Sharvari Deshpande, Carlyn Guthrie, Xuejuan Tan, Eliezer Aimontche, Tong Zheng, Andrew S. Greenberg, Xiang-Dong Wang, Koji Kitazawa, Gregory Dolnikowsi, Yong Cheng, Nirupa R. Matthan, Arvind Ramanthan, Jussi Pihlajamaki, Ron Korstanje, Christopher D. Wiley

## Abstract

When subject to damage or stress, cells develop responses in order to maintain tissue homeostasis. Two such decisions are ferroptosis and cellular senescence, but how cells decide between these outcomes remains unclear. Here we show that senescent cells increase levels of multiple membrane-bound polyunsaturated fatty acids (PUFAs), but a specific PUFA, dihomo-gamma-linolenic acid (DGLA, 20:3*ω*-3) is reduced. Exogenous repletion of DGLA or inhibition of delta-5-desaturase, the enzyme that metabolizes DGLA, instead results in cell death by ferroptosis. Senescent cells had elevated levels of other fezzroptosis sensitizers, including labile iron and expression of lipoxygenases – but also increased Gpx4 levels to prevent ferroptosis. Oral DGLA lowered senescent cell burden in aged mice and improved age-related functional outcomes. Finally, obese humans with lowered DGLA desaturation rates showed lower markers of adipose tissue senescence. Together, our data implicate DGLA and its desaturation as a major driver of decisions between senescence and ferroptosis.

## Introduction

Throughout the lifespan cells are subject to various insults and stressors, including macromolecular damage, epigenetic disruption, metabolic shifts, and mitochondrial dysfunction^1,2^. If unresolved, accumulation of these insults over time can result in degenerative conditions and cancer, necessitating activation of protective cellular responses. Two such responses are cellular senescence and ferroptosis. Senescence is a highly variable state primarily consisting of stable proliferative arrest coupled to the secretion of a complex mélange of biologically active molecules including cytokines, growth factors, proteases, and oxylipins that can have potent effects on tissue and systemic homeostasis^3–5^. Accumulation of senescent cells drives age-related pathology and limits lifespan, as methods that selectively kill senescent cells protect against age related disease and extend lifespan in murine models^6–8^. Ferroptosis is a form of lytic cell death that occurs through iron-dependent propagation of membrane-bound lipid peroxides^9^. It is hypothesized to be proinflammatory as lysis releases intracellular contents, including multiple immunogenic damage-associated molecular patterns (DAMPs), into the extracellular space^10^.

Senescence and ferroptosis share multiple features in common, as both are stress responses with tumor suppressive activities^11,12^. Indeed, multiple inducers of senescence, including oncogenic RAS, genotoxic stress, mitochondrial dysfunction, and activation of cell cycle arrest are known to sensitize cells to ferroptosis^9,13–15^. Despite these links, the relationship between senescence and ferroptosis remains relatively underexplored.

We previously identified oxylipin synthesis as a key feature of senescence that both reinforces the cell cycle arrest and promotes multiple segments of the senescence-associated secretory phenotype^3^. As polyunsaturated fatty acids (PUFAs) are the substrates used for synthesis of these oxylipins, we sought to determine if PUFA metabolism is altered in response to induction of senescence, and what the consequences of these alterations might be. From these studies, we identified alterations in PUFA metabolism that protect senescent cells from ferroptosis by lowering levels of the pro-ferroptotic PUFA, dihomo-gamma-linolenic acid (DGLA).

## Results

### Alterations in PUFA desaturation promote senescent cell survival

We previously observed increased synthesis of oxylipins in senescent cells^3^, as well as increased levels of many free PUFAs. Since membranes are a major source of PUFA for oxylipin biosynthesis, we sought to investigate PUFA levels in the cellular membrane. Following membrane isolation, fatty acids were extracted, methylated, and quantified using gas chromatography (GC). Induction of senescence increased the proportion of PUFA in membranes relative to saturated and monounsaturated fatty acids (SFA and MUFA) (**Figure S1A**), due to increased proportions of both ω-6 and ω-3 fatty acids (**Figure 1A**). While most individual ω-6 and ω-3 PUFA were increased in senescent cell membranes (**Figures 1B and 1C**), intermediates between the products of delta-6-desaturase (D6D, gene name FADS2) and delta-5-desaturase (D5D, gene name FADS1), specifically dihomo-gamma-linolenic acid (DGLA) (**Figure 1D**) and stearidonic acid (SDA) (**Figure 1E**) were notably decreased in senescent cells. Consistent with this, estimated D6D activity was decreased during senescence, whereas D5D activity was increased for both ω-6 (**Figure S1B**), and ω-3 (**Figure S1C**) fatty acids. Using our previously described mass spectrometry dataset^3^, we observed similar results for free fatty acids in senescent cells induced by either ionizing radiation [SEN(IR)] or mitochondrial dysfunction (MiDAS) (**Figure S1D**). Consequently, D5D:D6D desaturase activity ratios were highly elevated for both ω-6 (**Figure 1F**), and ω-3 (**Figure 1G**) fatty acids in membranes, as well as free ω-6 fatty acids (**Figure 1H**). Similar results were obtained when senescent cells were analyzed for phospatidylcholines (PCs) (**Figure S1E**), lysophosphatidylcholines (LPCs) (**Figure S1F**), and cholesterol esters (CEs) (**Figure S1G**), suggesting that it is desaturation imbalance that drives this effect. To confirm this, we measured *FADS1:FADS2* (which encode D5D and D6D, respectively) RNA ratios by quantitative PCR for multiple inducers of senescence, including ionization radiation (IR), RasV12 overexpression (RAS), treatment with nutlin-3a (Nut), and mitochondrial dysfunction (MiDAS) (**Figure 1I**). In each case, the expression of *FADS1* relative to *FADS2* was increased during senescence. Thus, induction of senescence is coupled to alterations in PUFA desaturation that reduce levels of intermediate PUFAs.

**Figure 1.**
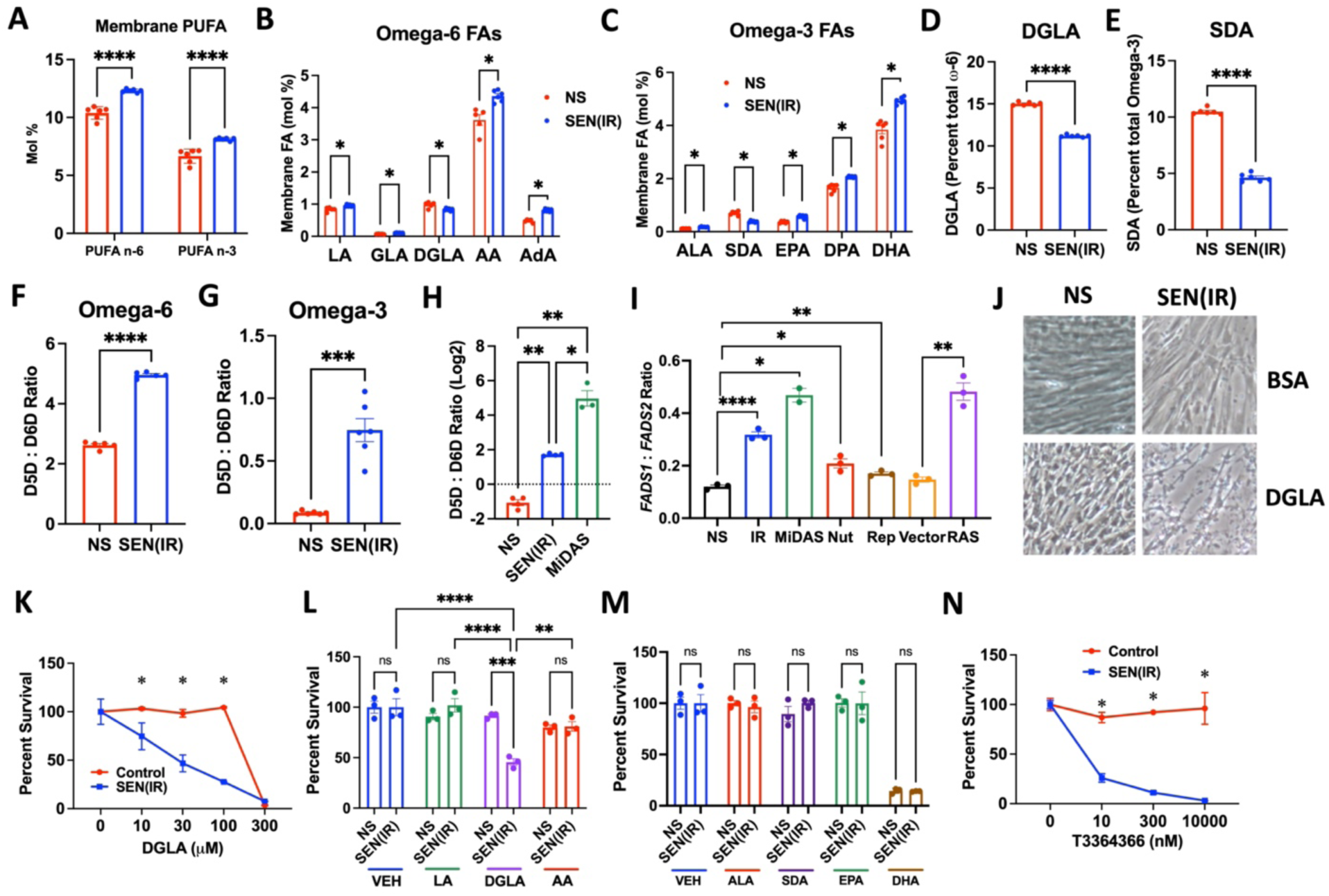
Altered PUFA desaturation promotes survival of senescent cells. A-G. Cellular membranes were extracted from IR-induced senescent [SEN(IR)] or mock-irradiated non-senescent (NS) cells 72 h after treatment and analyzed by GC for the indicated fatty acids. A. Relative levels of ω-6 and ω-3 PUFA in membranes of NS and SEN(IR) cells. B. Relative levels of indicated ω-6 PUFA (linoleic acid – LA, gamma-linolenic acid – GLA, dihomo-gamma-linolenic acid - DGLA, arachidonic acid – AA, adrenic acid – AdA). C. Relative levels of indicated ω-3 PUFA (alpha-linolenic acid - ALA, stearidonic acid - SDA, eicosapentanoic acid – EPA, docosapentanoic acid – DPA, docosahexanoic acid – DHA). D. DGLA percentage of ω-6 PUFA. E. SDA percentage of ω-3 PUFA. F. Ratios of D5D desaturation to D6D desaturation for ω-6 PUFA. G. Ratios of D5D desaturation to D6D desaturation for ω-3 PUFA. H. HPLC-MS data from Wiley et. al. ^3^ was analyzed for D5D desaturation (AA:DGLA) to D6D (DGLA:AA) desaturation ratios for ω-6 PUFA. I. Cells were induced to senesce by ionizing radiation (IR), antimycin A (MiDAS), nutlin-3a (Nut), serial culture to replicative exhaustion (Rep), and lentiviral RasV12 overexpression (RAS). Non-senescent (NS) and empty lentiviral vector (Vector) cells were used as controls. J. Representative images of NS or SEN(IR) cells treated with DGLA (200 μM) or carrier (BSA) for 6 days. K. Control (NS) or SEN(IR) cells treated with the indicated concentration of DGLA for 3 days, followed by measurement of survival by CCK8. L. NS or SEN(IR) cells were treated with 200 μM of each of the indicated ω-6 PUFA for 3 days, followed by measurement of survival by CCK8. M. NS or SEN(IR) cells were treated with 200 μM of each of the indicated ω-3 PUFA for 3 days, followed by measurement of survival by CCK8. N. Control (NS) or SEN(IR) cells were treated with the indicated concentration of T3364366 for 7 days, followed by measurement of survival by cell counting.

We next investigated the role of DGLA in senescence. We cultured non-senescent (NS) control or senescent [SEN(IR)] cells in the presence of DGLA (200 μM) or vehicle (BSA). Surprisingly, we observed that after 3 days senescent, but not non-senescent, cells cultured in the presence of DGLA underwent cell death (**Figure 1J**). DGLA was similarly toxic to replicatively senescent fibroblasts (**Figure S1H**). DGLA increased cell numbers in other non-senescent cell types, and killed them when induced to senesce - with hepatic stellate cells (HSteC) induced to senesce by oncogene activation or mitochondrial dysfunction (**Figure S1I**), mesenchymal stem cells (MSC) (**Figure S1J**), or HepG2 hepatocellular carcinoma cells treated with either nutlin-3a or palbociclib (**Figure S1K**), confirming broad cytotoxic activity of DGLA toward senescent cells. We confirmed that this killing was dose-dependent (**Figure 1K**), and similar results were observed for the DGLA precursor, gamma-linolenic acid (GLA) (**Figure S1L**). Furthermore, following treatment on NS and SEN(IR) cells with equimolar (200 μM) concentrations of other ω-6 (**Figure 1L**), and ω-3 (**Figure 1M**) fatty acids show no selective toxicity, confirming that the selective toxicity is DGLA-specific. These effects were not due to the combination of carbon chain length or number of double bonds, as dihomo-alpha-linolenic acid (DALA, 20:3 in the ω-3 conformation) (**Figure S1M**) and Mead’s acid (20:3 in the ω-9 conformation) (**Figure S1N**). Finally, blockade of DGLA desaturation using a D5D inhibitor (T3364366) specifically and dose-dependently resulted in death of senescent, but not non-senescent cells (**Figure 1L**). Inhibitors of other fatty acid metabolism pathways, including steroyl-CoA desaturase (SCD) (**Figure S1O**), fatty acid synthase (FASN) (**Figure S1P**), and D6D (**Figure S1Q**) had no selective toxicity to senescent cells. Together, these data suggest that desaturation imbalance between D6D and D5D promotes survival of senescent cells by lowering DGLA levels.

### DGLA kills senescent cells by ferroptosis

Since DGLA specifically and dose-dependently killed senescent cells, we sought to determine the mechanism of this cell death. DGLA is reported to kill cancer cells by both apoptosis^16^ and ferroptosis^17^. To differentiate between these possibilities, we treated either NS or SEN(IR) cells with DGLA, and measured survival (CytoCalcein violet – blue channel), apoptosis (Apopxin™ Green – green channel), and lytic cell death (7-amino-actinomycin D – red channel) (**Figure 2A**). Treating senescent cells with DGLA lowered cell viability (**Figure 2B**) and increased lytic cell death staining (**Figure 2C**). By comparison, while seeding on optical glass resulted in a basal level of apoptosis in all samples, these levels did not change upon DGLA administration (**Figure 2D**). Furthermore, treatment with 8-hydroxyoctanoic acid (8-HOA), a metabolic byproduct of DGLA metabolism that is reported to promote apoptosis^16^, had no selective effects on senescent cell survival, even at doses that exceeded DGLA treatments (**Figure S2A**).

**Figure 2.**
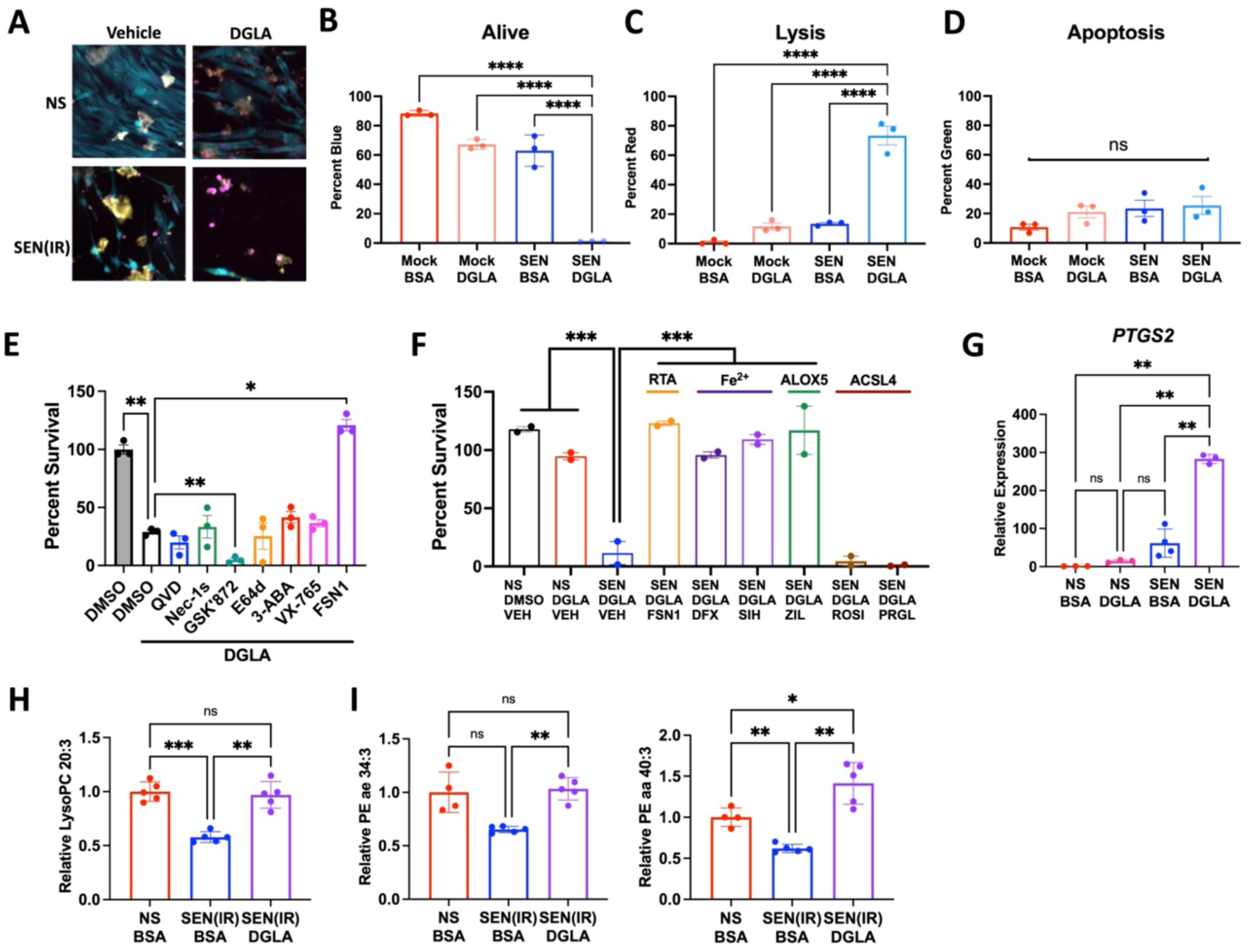
DGLA kills senescent cells by ferroptosis. A-D. NS or SEN(IR) cells were treated with either vehicle (BSA) or 200 μM DGLA for 48 hours, and were then stained for living cells, apoptotic cell, and lytic cells. A. Representative culture images for living (blue), lysis (red - pseudocolored violet), or apoptosis (green - pseudo-colored yellow). B. Percentage of each treatment group that remained alive. C. Percentage of each treatment group that underwent lysis. D. Percentage of each treatment group that underwent apoptosis. E. SEN(IR) cells were treated with either vehicle (BSA) or 100 μM DGLA for 72 hours in presence of inhibitors of various cell death pathways and cell survival was measured by CCK8 assay. F. NS or SEN(IR) cells were treated with either vehicle (BSA) or 100 μM DGLA for 72 hours in the presence of 5 μM ferrostatin-1 (FSN1), 50 μM deferoxamine (DFX), 30 μM salicylaldehyde isonicotinoyl hydrazine (SIH), 30 μM zileuton (Zil), 30 μM rosiglitazone (ROSI), or 100 μM PRGL493 (PRGL) and cell survival was measured by CCK8 assay. G. NS or SEN(IR) cells were treated with either vehicle (BSA) or 100 μM DGLA for 48 hours, RNA was extracted, and *PTGS2* levels were measured by quantitative PCR normalized to actin. H-J. NS or SEN(IR) (7 d) were treated with 100 μM DGLA for 48 hours, followed by extraction and analysis of esterified fatty acid species by mass spectrometry using a commercial kit (Biocrates). H. DGLA lysophosphatidylcholines were measured by targeted mass spectrometry. I. Relative peak area for PE ae 34:3. J. Relative peak area for PE aa 40:3.

To determine the mechanism behind DGLA-induced death of senescent cells, we treated SEN(IR) cells with DGLA and either a vehicle (DMSO) or inhibitors of various cell death modalities. Inhibitors of apoptosis (QVD-OPH), necroptosis (Nec-1s and GSK’872), lysosome-dependent cell death (E64d), parthanatos (3-aminobenzamide, 3-ABA), and pyroptosis (VX-785) had no effect on DGLA-mediated cell death, while an inhibitor of ferroptosis (ferrostatin-1, FSN1) allowed survival of senescent cells in the presence of DGLA (**Figure 2E**). Ferroptosis can be suppressed by multiple molecules, including radical trapping antioxidants (RTAs) like FSN1, iron chelators (deferoxamine – DFX or salicylaldehyde isonicotinoyl hydrazine – SIH), lipoxygenase inhibitors (zileuton - Zil, itself an RTA), and ACSL4 inhibitors (rosiglitazone – ROSI or PRGL493 - PRGL) (**Figure 2F**). While FSN1, iron chelators, and zileuton all prevented ferroptosis induced by DGLA, ACSL4 inhibitors did not rescue these effects (**Figure 2F**). ACSL4 is a major acyl-CoA synthase for AA and its promotion of ferroptosis^18^, but it is unclear which ACSLs are the primary acyl-CoA synthases for DGLA, so it remains possible that another ACSL is responsible for DGLA-promoted ferroptosis.

Both senescence and ferroptosis result in increases in expression of cyclooxygenase 2 (gene name *PTGS2*)^3,19^. We therefore measured *PTGS2* RNA levels by quantitative PCR. While senescent cells showed elevated levels of *PTGS2*, treatment with DGLA elevated these levels substantially (**Figure 2G**), consistent with induction of ferroptosis in senescent cells.

Since ferroptosis occurs at cellular membranes, we next sought to determine if treatment with DGLA resulted in repletion of DGLA in membrane phospholipids. Indeed, we observed increases in membrane-bound DGLA in control and senescent cells following DGLA administration (**Figure S2B**). Furthermore, mass spectrometry confirmed that administration of senolytic doses of DGLA (200 μM) restored DGLA-lysophosphatidylcholines levels (**Figure 2H**) – measured using targeted lipidomics (Biocrates). Since phosphatidylethanolamines (PE) on the inner leaflet are drivers of ferroptosis^20^, we used untargeted analysis of the same experiment to confirm that putative levels of PE ae 34:3 (**Figure 2I**) and PE aa 40:3 (**Figure 2J**) were increased by DGLA to at least non-senescent levels. While multiple combinations of fatty acids can result in these PE signatures, both were increased by DGLA administration, suggesting that DGLA incorporation drives this increase. We also observed elevated levels of several other esterified PUFA species following DGLA (**Figure S2C**). Together, these data confirm that repletion of DGLA selectively induces ferroptosis in senescent cells.

### Senescent cells are susceptible to ferroptosis

Since senescent cells died by ferroptosis when exposed to DGLA, we hypothesized that they might generally be more prone to ferroptosis than non-senescent cells. Three major drivers sensitize cells to ferroptosis: membrane-bound PUFA, intracellular labile iron, and an initiating lipid peroxidation event^21,22^. Since we observed increased membrane-bound PUFA in senescent cells (**Figure 1A**), we used a targeted lipidomics dataset (Biocrates) to measure multiple classes of esterified fatty acids (**Figure 3A**). Notably, senescent cells had elevated levels of PUFA phosphatidylcholines with esterified fatty acids, but reduced levels of PUFA phosphatidylcholines with ether bonds. Commensurate with an increase in membrane PUFA, we observed a decrease in lysophosphatidylcholines (LPCs) and PUFA diglycerides (DGs), with DG-O(16:0,20:4) being the most notably reduced species. In agreement with these findings, senescent cells had a reduced total ester to ether PC ratio (**Figure 3B**). Since ether phospholipids tend to be pro-ferroptotic in mammalian cells^23^, it is unclear if these alterations in PUFA composition are on balance more likely to promote or suppress ferroptosis in senescent cells.

**Figure 3.**
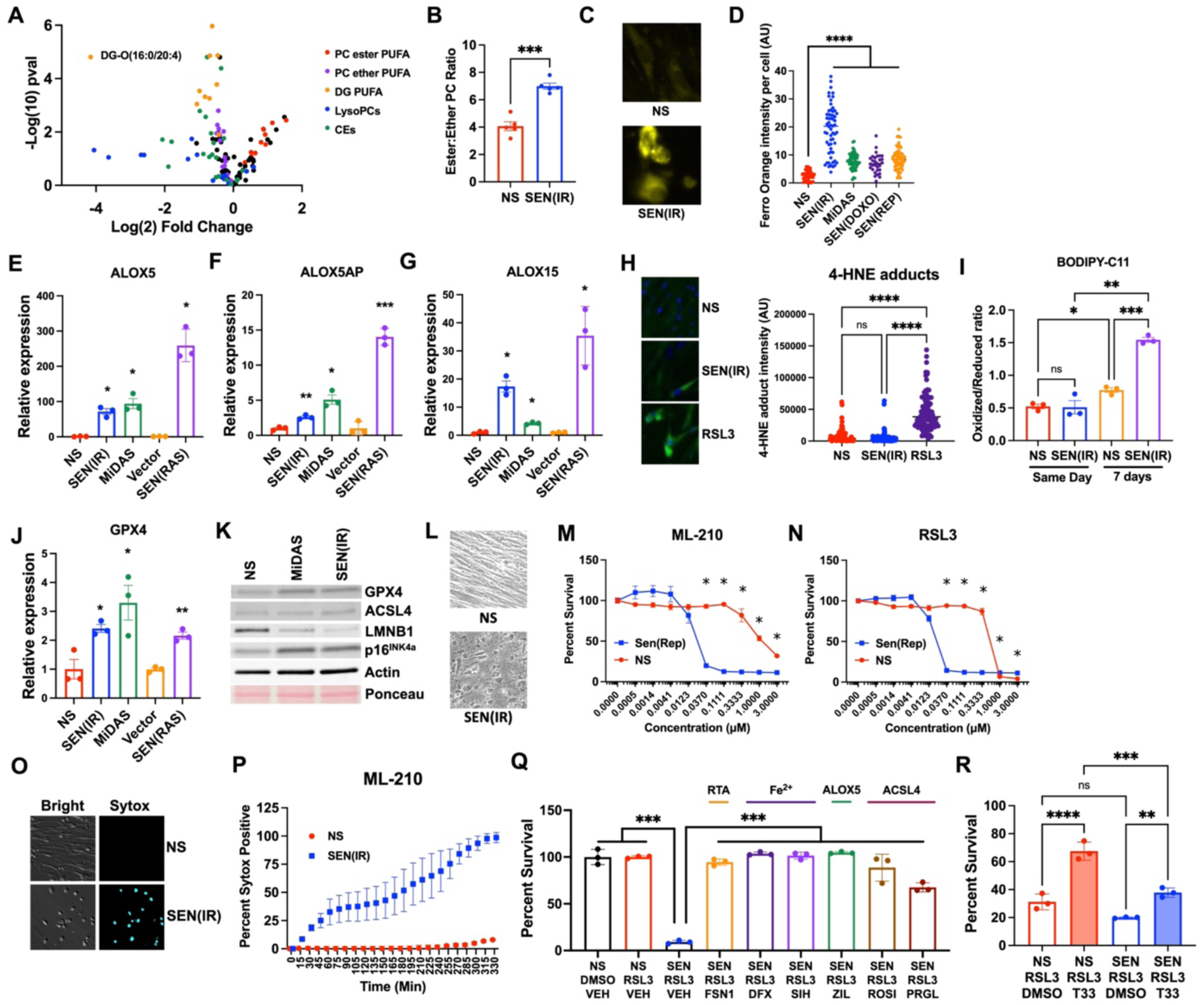
Senescent cells are sensitive to ferroptosis. A. Volcano plot of levels of esterified PUFA species in SEN(IR) relative to NS IMR-90 fibroblasts. B. Ratio of total ester to total ether phosphatidylcholines (PCs) in NS and SEN(IR) IMR-90 fibroblasts. C. Representative images of NS or SEN(IR) IMR-90 fibroblasts stained with FerroOrange D. IMR-90 fibroblasts were induced to senesce by 10 Gy IR [SEN(IR)], 250 nM antimycin A (MiDAS), 24 hours in 250 nM doxorubicin [SEN(DOXO)], or serial passage [SEN(REP)] and stained with FerroOrange on day 10 after induction and fluorescence intensities for at least 30 cells were quantified using ImageJ. E-G. IMR-90 fibroblasts were induced to senesce by IR, antimycin A, or overexpression of RasV12 as above and 10 days later RNAs were extracted and analyzed for the indicated transcripts by quantitative PCR, normalized to actin. E. RNA levels of *ALOX5*. F. RNA levels of *ALOX5AP*. G. RNA levels of *ALOX15*. H. (Left) Representative images of immunofluorescence (green) for 4-HNE adducts in NS, SEN(IR), or RSL3-treated IMR-90 fibroblasts, counterstained with DAPI. (Right) Quantitation of 4-HNE fluorescence intensity for at least 30 cells per condition. I. NS and SEN(IR) cells were treated with BODIPY-C11 and imaged on the same day or washed out and imaged 7 days later. Ratios represent oxidized (green channel) to reduced (red channel) J. RNAs levels of *GPX4*, normalized to actin. K. Protein lysates from NS, MiDAS, and SEN(IR) cells were analyzed for GPX4, ACSL4, LMNB1, p16^INK4a^, and beta-actin. Ponceau staining serves as a second loading control. L. Representative bright field images of NS or SEN(IR) cells treated with ML-210 (300 nM) for 3 hours. M. Sen(Rep) or NS (PD < 30) were treated with the indicated doses of ML-210 and were analyzed 18 hours later by CCK8. N. Sen(Rep) or NS (PD < 30) were treated with the indicated doses of RSL3 and were analyzed 18 hours later by CCK8. O. Representative bright field and Sytox blue images of NS or SEN(IR) cells treated with RSL3 (1 μM) for 3 hours. P. NS or SEN(IR) cells were given 1 μM ML-210 and percent Sytox positive cells (normalized to end point Hoechst) were calculated for 3 wells at the indicated timepoints. Q. NS or SEN(IR) cells were treated with 100 nM RSL3 and either vehicle FSN1, DFX, SIH, ZIL, ROSI, or PRGL – and survival was measured by CCK8 assay. Data are expressed as percent NS or SEN(IR) treated with DMSO. R. NS or SEN(IR) HepG2 cells were treated with 100 nM RSL3 in the presence or absence of 10 μM T3364366 for 18 hours and cell survival was measured by CCK8 assay normalized to DMSO-treated NS or SEN(IR) cells.

Labile iron catalyzes the Fenton reaction that propagates lipid peroxidation and results in ferroptosis^9^. In agreement with previous work^24–26^, we observed increased labile iron in senescent cells, as measured by FerroOrange staining (**Figure 3C**), regardless of inducer (**Figure 3D**) or cell type (**Figure S3A**). Additionally, we previously showed that senescent cells elevate synthesis of multiple oxylipin synthases^3,27,28^, and many of these, such as the arachidonate lipoxygenases (ALOXs) can catalyze peroxidation of plasma membrane phospholipids^29^. In agreement with our previous work, we confirmed that senescent cells elevate expression of *ALOX5*, *ALOX5AP*, and *ALOX15* in response to multiple inducers (**Figures 3E-3G**). Finally, we measured lipid peroxidation via immunofluorescence staining for 4-HNE adducts (**Figure 3H**) and BODIPY-C11 oxidized/reduced ratios (**Figure 3I**). While 4-HNE and early BODIPY-C11 staining did not indicate any increased lipid peroxidation in senescent cells, BODIPY-C11 became slightly more oxidized over the course of 7 days in control cells, and extensively oxidized in senescent cells (**Figure 3I**). Together, these data suggested that senescent cells have features that might sensitize them to ferroptosis, but are somehow protected from rapid lipid peroxidation and cell death.

Ferroptosis is antagonized by multiple factors, most notably by the activity of glutathione peroxidase 4 (Gpx4), which catalyzes the glutathione-dependent conversion of fatty acid peroxides to fatty acid hydroxyls^19,30^. Quantitative PCR analysis revealed increases in *GPX4* RNA levels in senescent cells induced by IR, mitochondrial dysfunction, or oncogene activation (**Figure 3J**). Western blots similarly revealed that both SEN(IR) and MiDAS cells had elevated GPX4 protein levels, but no change in ACSL4 (**Figure 3K**), suggesting that GPX4 might protect senescent cells from ferroptosis, but that altered accumulation of PUFA during senescence is not due to increases in ACSL4 levels. We therefore tested the Gpx4 inhibitors ML-210 and RSL3 for the ability to selectively induce ferroptosis in senescent cells. Within hours of treatment with ML-210 - SEN(IR), but not NS cells - began to display morphological features of cell death (**Figure 3L**) and after 24 hours showed dose-dependent selective killing (**Figure S3B**). Both ML-210 and RSL3 selectively killed SEN(REP) cells (**Figures 3M-3N**). RSL3 was also selectively toxic to doxorubicin-induced senescent cells [SEN(DOXO)] (**Figure S3C**), but the ferroptosis inducer erastin, which induces ferroptosis by glutathione depletion^19^, was less effective (**Figure S3D**). Similar results were observed with senescent astrocytes (**Figure S3E-S3F**) and HepG2 cells (**Figure S3G-S3H**). We further confirmed the kinetics of ferroptosis following GPX4 inhibition using Sytox staining (**Figure 3O**) and observed more rapid cell killing in senescent cells relative to non-senescent in response to ML-210 (**Figure 3P**), RSL3 (**Figure S3I-S3J**), or FIN-56 (**Figure 3K**). Much like DGLA, ferroptosis induced by Gpx4 inhibition was blocked by FSN1, iron chelators, and zileuton (**Figure 3Q**), but unlike DGLA, pre-treatment with ACSL4 inhibitors prevented ferroptosis. Surprisingly, short term inhibition of D5D/FADS1 with T3364366 antagonized ferroptosis induced by GPX4 inhibitors (**Figure 3R**), consistent with previous reports that FADS1 depletion protects against ferroptosis following GPX4 inhibition^31,32^. Together, these data indicate that DGLA-induced ferroptosis of senescent cells follows a pattern that shares features with, but is mechanistically distinct from, GPX4 inhibitor-induced ferroptosis.

### DGLA lowers senescent cell burden and improves aging phenotypes in mice

Since DGLA selectively induced ferroptosis in senescent cells, we sought to determine if it could perform similarly *in vivo*. We therefore gavaged aged (22-24 mo) p16-3MR (**Figure 4A-4C**) or C57Bl/6J (**Figure 4D-4X**) mice with either DGLA ethyl ester (DGLA-EE) or vehicle (phosal-50) for five consecutive days, followed by a two week recovery period. We selected DGLA-EE for *in vivo* bioavailability purposes but confirmed the selective killing of senescent cells using DGLA-EE in culture (**Figure S4A**). Treatment of aged p16-3MR mice with DGLA lowered whole body luminescence, a proxy for p16 expression (**Figure 4A-4B**). Furthermore, fat and liver tissues isolated from DGLA-treated mice showed lower senescence-associated beta-galactosidase staining than those from vehicle-treated mice (**Figure 4C**). RNA from adipose tissue from DGLA-treated aged C57Bl/6J mice showed lower levels of several SASP factors, as did livers from aged male, but not female mice (**Figure 4D**). Adipose tissue RNA levels of several cyclin-dependent kinase inhibitors, including *p16^INK4a^* (**Figure 4E**), *p15^INK4b^* (**Figure 4F**), and *p21^WAF1^* (**Figure 4G**) were similarly lowered by DGLA administration. Thus, DGLA lowers senescent cell burden in aged animals.

**Figure 4.**
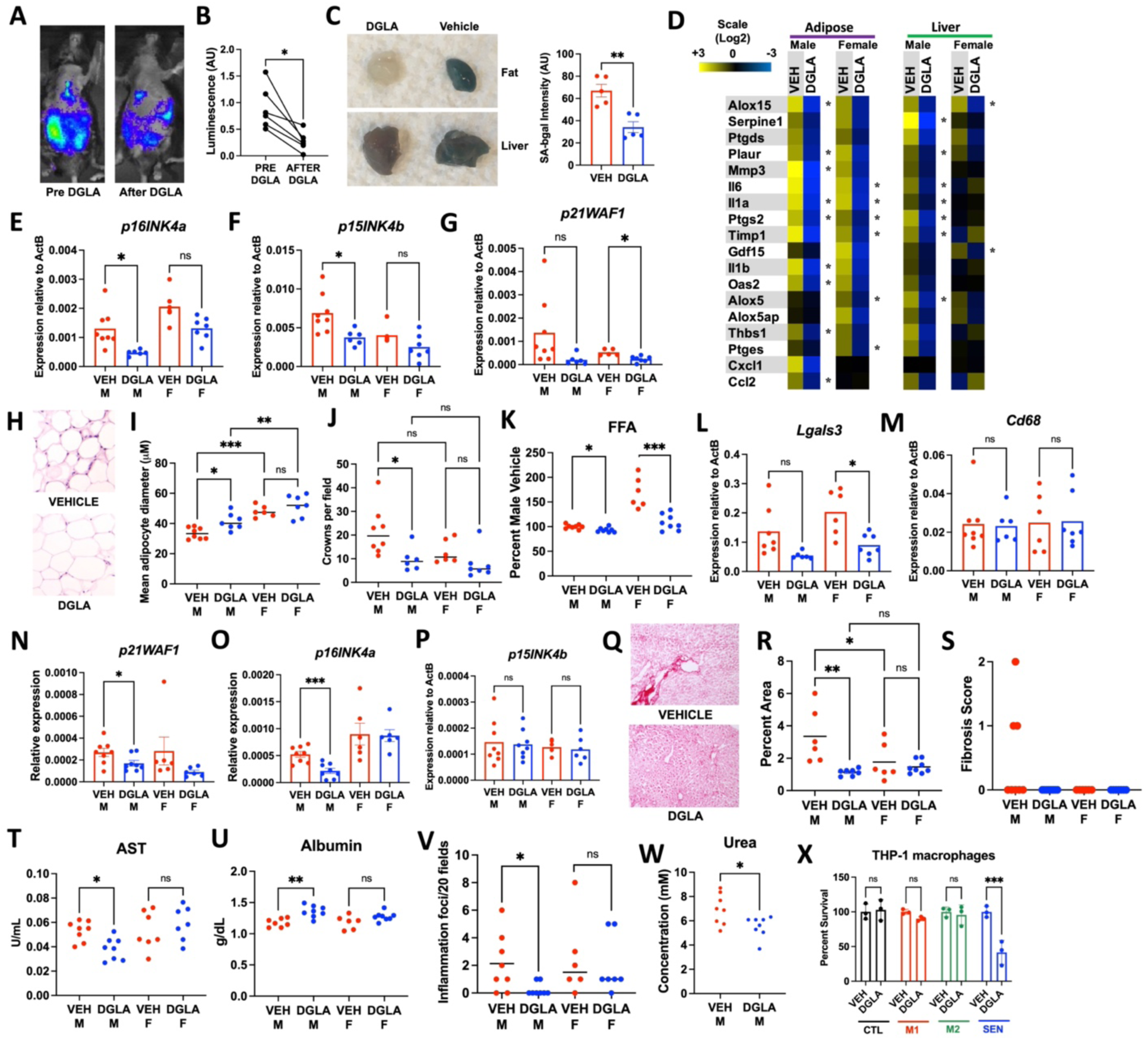
DGLA kills senescent cells and improves aging phenotypes in mice. Aged (22-24 months) p16-3MR (A-C) or C57Bl/6J (D-X) mice were gavaged with 400 mg/kg DGLA-EE in Phosal-50 for 5 consecutive days and allowed to recover for 2 weeks before analysis and tissue harvest. P16-3MR mice were shaved before luminescence imaging. A. Representative luminescence images of the same p16-3MR mouse before and after DGLA-EE treatment. B. Quantitation of luminescence levels of 6 aged mice before and after treatment with DGLA-EE, normalized to readings from the same phantom. *= p<0.05, paired, 2-tailed t-test. C. (Left) representative images of senescence-associated beta-galactosidase in fat and liver from aged mice treated with either DGLA-EE or vehicle (phosal-50). (Right) blue intensity was quantified with ImageJ. D. SASP factor RNA expression was measured by quantitative PCR in adipose and liver tissues from aged male and female mice. E. RNA from adipose tissue from male and female mice treated with DGLA-EE or vehicle was analyzed by qPCR for *p16^INK4a^*. F. RNA from adipose tissue from male and female mice treated with DGLA-EE or vehicle was analyzed by qPCR for *p15^INK4b^*. G. RNA from adipose tissue from male and female mice treated with DGLA-EE or vehicle was analyzed by qPCR for *p21^WAF1^*. H. Representative image of H&E stained adipose tissue from vehicle or DGLA-EE treated mice. I. Mean adipocyte diameters were calculated from 3 fields per animal using Adiposoft. Each point represents the mean adipocyte diameter of one animal. J. Crown-like structures were quantified per field using the means of 3 fields per animal. Each point represents a single animal. K. Free fatty acids (FFA) were measured in blood sera from vehicle and DGLA-EE treated mice. L. RNA from adipose tissue from male and female mice treated with DGLA-EE or vehicle was analyzed by qPCR for *Lgals3*. M. RNA from adipose tissue from male and female mice treated with DGLA-EE or vehicle was analyzed by qPCR for *Cd68*. N. RNA from liver tissue from male and female mice treated with DGLA-EE or vehicle was analyzed by qPCR for *p21^WAF1^*. O. RNA from liver tissue from male and female mice treated with DGLA-EE or vehicle was analyzed by qPCR for *p16^INK4a^*. P. RNA from liver tissue from male and female mice treated with DGLA-EE or vehicle was analyzed by qPCR for *p15^INK4b^*. Q. Representative images of picrosirius red stained frozen liver sections from vehicle or DGLA-EE treated male mice. R. Picrosirius red positive fibrotic area was calculated using 3 fields from vehicle and DGLA-EE treated male and female mice using ImageJ. Each point represents the averages of 3 fields per animal. S. Fibrosis pathology scores for livers of mice treated with vehicle or DGLA-EE. T. Aspartate aminotransferase (AST) activity in blood sera from mice treated with vehicle or DGLA-EE. U. Albumin levels in blood sera from mice treated with vehicle or DGLA-EE. V. Histological counts of age-associated inflammation foci in 20 fields per animal. W. Urea levels in blood sera from mice treated with vehicle or DGLA-EE. X. THP-1 macrophages were induced into M1 or M2 polarization, or induced to senesce by doxorubicin, and then treated with DGLA for 72 hours, followed byt

Senescent cells have been shown to drive age-related lipoatrophy and inflammation in adipose tissue. Histological analyses of adipose tissue sections from DGLA-treated mice revealed notable improvements (**Figure 4H**), including increases in adipocyte diameter (**Figure 4I**) and decreases in adipocytes per field (**Figure S4B**) and crown-like structures (**Figure 4J**). Free fatty acids (FFA), a marker of lipolysis, were similarly reduced in sera from DGLA-treated animals, with both a larger amplitude and larger reductions in female mice (**Figure 4K**). In addition, galectin-3 (*Lgals3*), a marker of macrophage activation – but also a SASP factor^33^ - was reduced by DGLA (**Figure 4L**), while the monocyte marker *Cd68* was unchanged (**Figure 4M**). Thus, elimination of senescent cells by DGLA is linked to reductions in inflammation and lipolysis.

We next evaluated the effects of DGLA on liver senescence, histology, and function. While DGLA lowered RNA levels of *p21^WAF1^* (**Figure 4N**) in both sexes, only males were statistically significant and *p16^INK4a^* levels were only lowered by DGLA in males (**Figure 4O**). By comparison, *p15^INK4b^* levels were unchanged by treatment (**Figure 4P**). Aged male mice showed markers of fibrosis as determined by picrosirius red staining (**Figure 4Q**) and confirmed by Masson’s trichrome, while DGLA-treated males showed less fibrotic area, and females showed little to no fibrotic staining (**Figure 4R**). Fibrosis pathology scores confirmed mild to moderate fibrosis for three vehicle-treated aged males, and no pathology for any other group (**Figure 4S**). DGLA, however, increased steatotic area in treated mice (**Figure S4C**), possibly in order to store the excess administered DGLA. Functionally, DGLA lowered serum AST levels in aged male, but not female mice (**Figure 4T**). Conversely, DGLA elevated serum albumin levels in aged male, but not female mice (**Figure 4U**). An additional marker of liver aging is the presence of inflammation foci^34^, and these foci were reduced by DGLA in male mice (**Figure 4V**). Analogous to adipose tissue, *Lgals3* expression was decreased in DGLA-treated male mice (**Figure S4D**), but *Cd68* remained unchanged (**Figure S4E**). Thus, liver showed a sexually dimorphic response to DGLA in terms of senescent cell clearance and functional improvement. Finally, DGLA lowered serum urea levels, which is inversely associated with kidney function, in aged male mice (**Figure 4W**). As such, these data indicate that DGLA-mediated elimination of senescent cells can improve multiple aging parameters *in vivo*.

Since we observed lowering of multiple markers of inflammation in both adipose and liver tissues - and activated macrophages can acquire multiple features of senescence^35–37^ - we considered the possibility that DGLA administration might kill or inactivate macrophages rather than kill senescent cells. We therefore induced M1 or M2 polarization, or treated macrophages with the senescence-inducing agent doxorubicin and assessed cell survival by CCK8 assay or markers of activation by quantitative PCR, followed by treatment with DGLA. DGLA had no effect on the survival of control, M1, or M2 polarized human THP-1 (**Figure 4X**) or murine RAW267.4 (**Figure S4F**) macrophages, but selectively killed both cell types following doxorubicin treatment. DGLA also increased *LGALS3* in M1 human macrophages (**Figure S4G**), rather than the decrease observed in DGLA-treated adipose and liver tissues. DGLA had no significant effect on two additional markers of macrophage activation, *ARG1* or *IL10* (**Figures S4H-S4I**), but did lower TNF-a levels in both M1 and M2 macrophages (**Figure S4J**). Furthermore, *CD38* can be induced in macrophages by the secretions of senescent cells^38,39^. *CD38* was elevated in M1 macrophages and reduced by DGLA (**Figure S4K**). However, no changes in *Cd38* were observed in the adipose tissue of DGLA-treated animals (**Figure S4L**). Thus, the effects of DGLA on aged mice are unlikely to be due to elimination of macrophages or suppression their activation.

### DGLA desaturation positively associates with senescence markers in human obesity

Human genetic variations at the FADS1 locus are a major determinant of DGLA desaturation^40^. In particular, the FADS1 single nucleotide polymorphism rs174547 has been highly studied in the context of PUFA desaturation and human disease, with the CC genotype associated with lowered DGLA desaturation rates and the TT genotype indicative of higher DGLA desaturation^40^. In agreement with this, the CC genotype is associated with reduced rates of stroke and cardiovascular disease^41,42^, metabolic syndrome^43^, and age-related macular degeneration^44^, phenotypes in which senescent cells are also implicated^45–49^. Obesity is associated with accumulation of senescent cells, especially in the fat^50–52^. We therefore analyzed subcutaneous adipose tissue samples from the Kuopio Obesity Surgery (KOBS) study, a prospective observational study of the metabolic consequences of obesity surgery^53^. These samples were previously analyzed for rs174547 genotype^54^ and fatty acid composition in various tissues^55^. We measured adipose *P21^WAF1^*, *P16^INK4A^*, and *P15^INK4B^* RNA expression by quantitative PCR, and examined associations with genotype (**Figures 5A-5C**) or DGLA desaturation (**Figures 5D-5E**).

**Figure 5.**
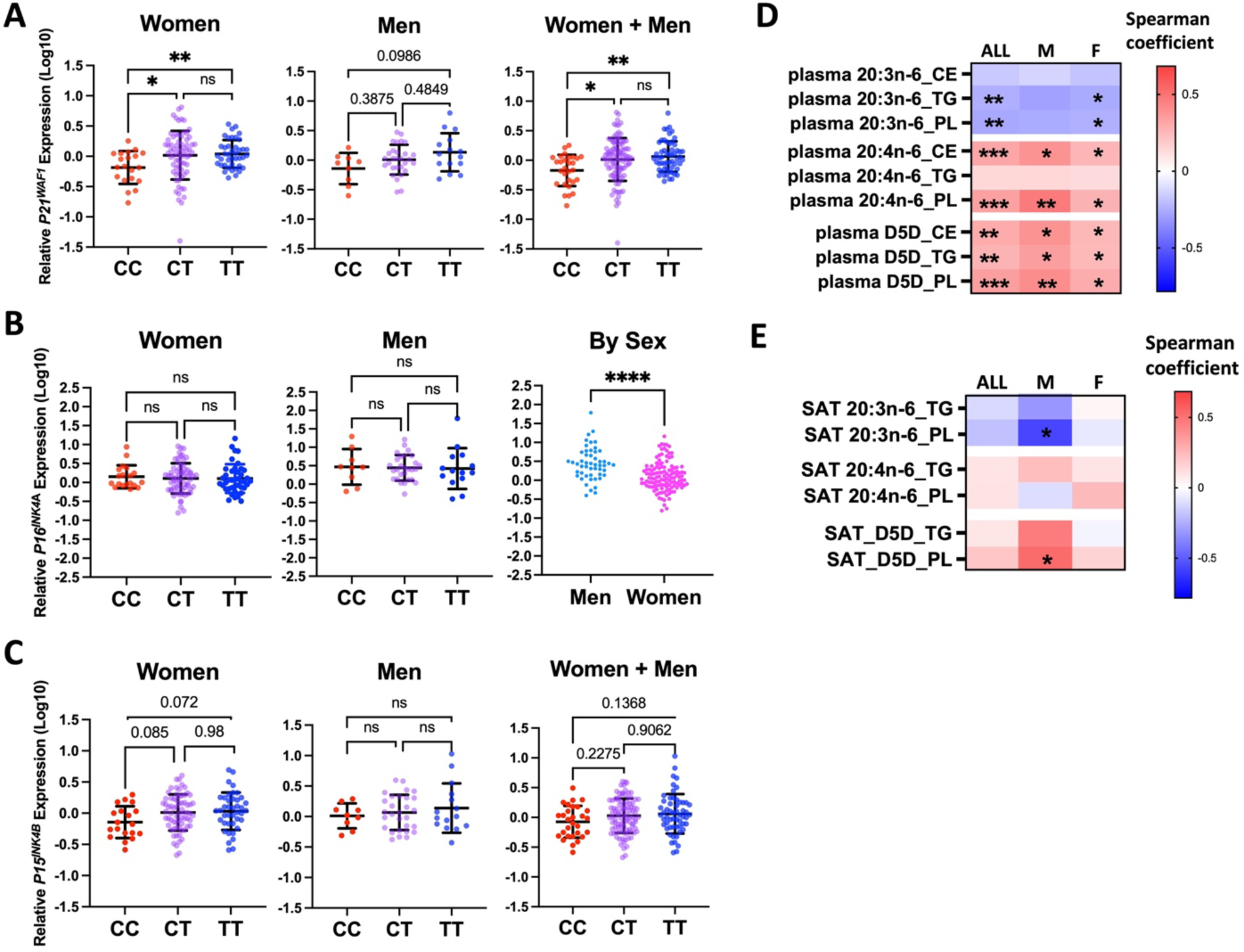
FADS1 activity is positively associated with cellular senescence in obese humans. **A-C**. Adipose tissue from KOBS study participants was genotyped for rs174547 and *P21^WAF1^*, *P16^INK4a^*, or *P15^INK4b^* normalized to beta actin. N for each group: Men = 54 total (10 CC, 29 CT, 15 TT). Women = 131 total (20 CC, 70 CT, 41 TT). **A.** Adipose *P21^WAF1^* RNA levels in women, men, and in combination by the genotypes of rs174547. (*=p<0.05, **=p<0.01, log-transformed Kruskal-Wallis test, Dunnett’s T3 adjustment for multiple comparisons). **B.** Adipose *P16^INK4a^* RNA levels in women (left), men (middle), by rs174547 genotype, or by sex (right). (****=p<0.0001, log-transformed Kruskal-Wallis test, Dunnett’s T3 adjustment for multiple comparisons). **C.** Adipose *P15^INK4b^* RNA levels in women, men, and in combination by rs174547 genotype. (Log-transformed Kruskal-Wallis test, Dunnett’s T3 adjustment for multiple comparisons). **D.** Spearman correlation coefficients for *P21^WAF1^* RNA levels and plasma levels of DGLA (20:3n-6), AA (20:4n-6), and D5D activity (AA:DGLA ratio) in cholesterol ester (CE), triglyceride (TG) and phospholipid (PL) fractions in women, men, and in combination. (*=p<0.05, **=p<0.01, ***=p<0.001, Spearman rank-order correlation coefficient test). **E.** Spearman correlation coefficients for *P21^WAF1^* RNA levels and subcutaneous adipose tissue levels of DGLA (20:3n-6), AA (20:4n-6), and D5D activity (AA:DGLA ratio) in triglyceride (TG) and phospholipid (PL) fractions in women, men, and in combination. (*=p<0.05, **=p<0.01, ***=p<0.001, Spearman rank-order correlation coefficient test).

We observed statistically significant decreases in *P21^WAF1^* RNA levels in women with the CC genotype compared to the CT and TT genotypes (**Figure 5A, left panel**). Men demonstrated similar trends in *P21^WAF1^* expression (**Figure 5A, middle panel**), but power was not sufficient to indicate significance (p = 0.099, CC to TT). By comparison *P16^INK4A^* levels did not vary with genotype (**Figure 5B)**, but were elevated in men relative to women (**Figure 5B, right panel**). Furthermore, we observed non-statistically significant (p = 0.072, CC to TT) trends in which the CC genotype was associated with reductions in *P15^INK4B^* in women, but less so in men. When women and men were combined, RNA levels of *P21^WAF1^* (**Figure 5A, right panel**), but not *P16^INK4A^* or *P15^INK4B^*, were significantly lower in the CC genotype. Thus, the rs174547 CC genotype is associated with reduced markers of adipose tissue senescence in humans, particularly *P21^WAF1^*.

We further sought to validate the relationship between DGLA desaturation and senescence by analyzing our previously described lipid profiling data^54,55^ for plasma (**Figure 5C**) and adipose tissue (**Figure 5D**) to calculate Spearman correlation coefficients between DGLA, AA, or D5D desaturation rates (AA:DGLA ratios) and *P21^WAF1^* expression, regardless of genotype. Plasma DGLA levels, regardless of species (CE, TG, or PL), were universally negatively correlated with *P21^WAF1^* levels, and AA levels were positively correlated with *P21^WAF1^* (**Figure 5C**). Furthermore, D5D activity was universally positively correlated with *P21^WAF1^*, and was statistically significant in each case. Similar but less robust results were observed in adipose tissue (**Figure 5D**), with men showing stronger correlations between *P21^WAF1^* and DGLA or D5D activity than women. In all cases, DGLA desaturation rates in phospholipids were most strongly associated with senescence, consistent with ferroptosis as a mechanism of elimination^53,56^. Thus, DGLA desaturation is associated with increased adipose tissue senescence in a cohort of humans with obesity. Together with our observations in culture and in mice, these data suggest that DGLA and its desaturation are a major determinant of senescent cell accumulation.

## Discussion

Cellular responses to stress and damage can occur via a variety of mechanisms, but how a cell decides between mechanisms such as senescence and ferroptosis remains unclear. The commonalities between inducers of senescence and sensitizers to ferroptosis suggest a link between these processes^9,13–15^. Now we show that PUFA desaturation imbalance is a key determinant that favors cellular senescence over ferroptosis. This imbalance lowers levels of membrane-bound DGLA and maintains survival of senescent cells. These results reveal a key pathway in the decision to adopt senescence and offer therapeutic avenues for intervention.

DGLA, but not other PUFA, selectively induced ferroptosis in senescent cells. This is somewhat surprising, as DGLA has fewer double bonds than AA and AdA that can be targeted for hydrogen abstraction and peroxidation^57^, and inhibition of D5D antagonizes short term induction of ferroptosis by GPX4 inhibitors (**Figure 3R**). This suggests that DGLA may drive ferroptosis by a GPX4-independent mechanism, most likely by being the preferred substrate of a ferroptosis-promoting enzyme. Previous work identified the epoxygenase activity of a cytochrome p450 enzyme as a primary driver of DGLA-mediated ferroptosis in nematodes^58,59^, suggesting a potential mechanism of action for future study.

Since mammals cannot synthesize PUFA *de novo*, they must acquire them from exogenous (dietary) sources^60,61^. PUFA levels in serially cultured cells are much lower than those found in freshly isolated cells or *in vivo*, at least in part due to culture of most cells in 10% fetal bovine serum, which itself has relatively low levels of PUFA^60^. As such, cells in an *in vivo* setting may undergo ferroptosis more frequently than in culture. However, thus far ferroptosis has primarily been studied as a driver of pathology, primarily due to embryonic lethality and tissue dysfunction that arise in constitutive and tissue-specific knockouts of Gpx4^30,61–63^. Here, we show multiple benefits from DGLA-mediated ferroptosis of senescent cells in aged animals. While the idea that ferroptosis is contextually beneficial outside of tumor suppression might seem surprising, Gpx4 knockout heterozygotes are long-lived^64^, suggesting the possibility that these animals benefit from conditions that might favor ferroptosis over senescence - though neither senescence nor ferroptosis has been measured in these animals.

To our knowledge, rs174547 represents the first human variant that might influence senescent cell accumulation via the decision to instead engage a cell death pathway. However, currently the relationship remains associative. Our data demonstrating the pro-ferroptotic effect of DGLA on senescent cells in culture and in mice strongly suggest that these effects are driven by ferroptosis of senescent cells in human tissue, but other possibilities remain. For example, we and others have shown that oxylipin derivatives of AA and AdA can reinforce senescence ^3,65,66^ – so it remains possible that lowered levels of DGLA desaturation prevent the development of senescence via reduction of AA, rather than elimination of senescent cells by ferroptosis.

Together, our data suggest a potential therapeutic opportunity for the elimination of senescent cells by targeting DGLA desaturation. By converting senescence into ferroptosis, elimination of senescent cells could allow for development of novel senolytic targets with therapeutic potential for multiple conditions driven by senescence. Similarly, these results might present an opportunity for cancer therapy by first inducing senescence in cancer cells and then driving their elimination by ferroptosis – two tumor-suppressive strategies that could be combined, as previously described for apoptosis^67,68^. Conversely, since ferroptosis can also drive tissue dysfunction, converting ferroptosis-promoting acute stressors into senescence instead might offer short term protection from tissue damage, as recently described for parthanatos^69^. If necessary, this could be followed by later elimination of senescent cells once tissue integrity is restored. Thus, identification of mechanisms by which cells decide between these cell fates is a potentially important first step in developing new therapies for cancer and degenerative diseases.

## Materials and Methods

### RESOURCE AVAILABILITY

#### Lead contact

Further information and requests for resources and reagents should be directed to and will be fulfilled by the Lead Contact, Christopher D. Wiley.

#### Material Availability

This study did not generate unique models or reagents.

### EXPERIMENTAL MODEL AND SUBJECT DETAILS

#### Cell lines and strains

Unless otherwise stated, all cells were IMR-90 fibroblasts cultured at 3% O2, as described in Wiley et. al., 2019^5^. IMR-90 fibroblasts (ATCC and Coriell), HepG2 (ATCC), bone marrow-derived mesenchymal stem cells (MSCs – a gift from Judith Campisi), HSteCs (ScienCell), and mouse RAW 264.7 macrophages (ATCC) were cultured in cultured Dulbecco’s Modified Eagle Medium (DMEM) with L-glutamine and 4.5g/L glucose, without sodium pyruvate and 10% fetal bovine serum (FBS) and 100 U/mL penicillin/streptomycin at 37°C, 5% CO2. **Human THP-1 macrophages were cultured at 37⁰C and 5% CO_2_ in RPMI 1640 medium supplemented with 10% and 100 U/ml streptomycin and penicillin, as previously described^70^.** Mouse-derived primary astrocytes were cultured in Dulbecco’s modified Eagle’s medium, N2 supplement (Invitrogen, #17502048), antibiotic/antmycotic (Invitrogen, #15240062), and supplemented with 35 µg/mL bovine pituitary extract (Invitrogen, #13028014), 5% FBS, 40 ng/mL epidermal growth factor (Sigma, #E9644), and 20 ng/mL bFGF (Sigma, #F0291). Astrocytes and macrophages were cultured in atmospheric O_2_, HSteCs and HepG2 were cultured in 6% O_2_ (which approximates liver oxygen levels), and fibroblasts and MSCs were cultured in 3% O_2_. All cells were confirmed mycoplasma free prior to use.

#### Animal models

P16-3MR experiments were conducted using a protocol (A10090) approved by the Institutional Animal Care and Use Committee of the Buck Institute. Mice were housed in alternating 12:12 hour light/dark cycles at 68-72°F and animal health rounds were performed on a daily basis, 7 days a week by Vivarium staff. Any illness or deaths were reported to the Vivarium Director and/or the Attending Veterinarian. The Buck Institute vivarium is an SPF barrier facility and health screening is done on all colonies on a facility-wide basis. Sentinel mice were screened quarterly after exposure to soiled bedding and byproducts from mice on the same rack for common murine viruses. Internal and external parasitology screening was also performed on all sentinel mice. P16-3MR mice were bred and maintained until the age of 22-24 months, imaged for luminescence, and then administered DGLA-EE (400 mg/kg) or vehicle (Phosal-50) for 5 consecutive days, and allowed to recover for 2 weeks before a second round of luminescence imaging.

#### Human Studies: KOBS study description

The present analysis also includes a validation cohort with cross-sectional baseline data from a sub-cohort of individuals with obesity (women: n=131, men: n=54) participating in the Kuopio Obesity Surgery (KOBS) study ^53^. Subjects with gene expression data from SAT and plasma FA composition available were included in analyses. The study protocol was approved by the Ethics Committee of the Northern Savo Hospital District (54/2005, 104/2008, and 27/2010, and 1108/2018) and carried out in accordance with the Helsinki Declaration. Written and oral information was given to the participants, and informed written consent was obtained from all participants. **FADS1 genotyping.** The study participants from the KOBS study were genotyped for the *FADS1* variant rs174547 using the TaqMan SNP Genotyping Assay according to the protocol provided by the manufacturer (Applied Biosystems, Foster City, CA, USA). **Fatty acid composition.** The analysis of FA composition in plasma and SAT lipid fractions was described previously ^55^. Briefly, plasma samples and pulverized SAT (40 mg) were extracted with chloroform–methanol (2:1), and the different lipid fractions, cholesteryl esters (CE), triglycerides (TG), and phospholipids (PL) were separated by solid phase extraction with an aminopropyl column. Samples were transmethylated with 14% boron trifluoride in methanol and were analyzed by 7890A gas chromatograph (Agilent Technologies, Inc., Wilmington, DE, USA) equipped with a 25-m FFAP column. Cholesteryl nonadecanoate (Nu Chek Prep, Inc., Elysian, MA, USA), trinonadecanoin, and phosphatidylcholine dinonadecanoyl (Larodan Fine Chemicals, Malmö, Sweden) served as internal standards. Enzyme activities in different lipid fractions were estimated as product-to-precursor ratios of individual FAs as follows: D5D = 20:4n-6 (AA)/20:3n-6 (DGLA). **Statistical analyses.** Genetic associations with continuous variables were analyzed with log-transformed Kruskal-Wallis tests using Dunnett’s T3 multiple comparisons test. The correlations between variables were analyzed using Spearman rank-order correlation coefficient tests.

### METHOD DETAILS

#### Induction of senescence

Sen(IR) cells were generated by exposing cells to 10 Gy of ionizing radiation or sham irradiation, as described previously^5^. MiDAS cells were generated by culturing with 500 nM antimycin A for seven days; or by depletion of mitochondrial DNA by culture in 100 ng/mL ethidium bromide until cell cycle arrest, as previously described^71^. SEN(RAS) was induced via transduction with a RasV12 lentivirus and empty vector served as a negative control. Cells were treated with lentivirus for 24 hours, after which the virus was removed and selected in puromycin for 48-72 hours and cells were used 7 days or more following the addition of Ras. Drug-induced senescent cells were treated with 5 mM of palbociclib or nutlin-3a for at least 5 days. SEN(DOXO) cells were treated with 250 nM doxorubicin for 24 hours, followed by return to growth media. Replicative senescent SEN(REP) cells were passaged until they stopped dividing (less than 5% EdU incorporation in 24 h), which occurred for IMR-90 fibroblasts at ∼70 population doublings.

#### Senescence-associated beta-galactosidase

Senescence-associated beta-galactosidase staining was conducted for both tissue and cultured cells using the Cell Signaling Technology Senescence β-Galactosidase Staining Kit (9860), following manufacturer instructions. Cells were fixed for 10 min, and tissues were fixed for 15 min. Staining intensity for tissue was calculated from captured images using ImageJ.

#### CCK-8 Viability Assays

Cells in cell-death assays were quantified using Cell Counting Kit-8 (CCK8, Dojindo, CK04), according to manufacturer protocol. Absorbance was measured on the Varioskan LUX Multimode Microplate Reader.

#### Macrophage Viability Assays

For mouse M1 macrophages, RAW 264.7 cells were pretreated with 20 ng/ml recombinant mouse IFN-γ (R&D Systems, #NBP2-76178) and 100 ng/ml lipopolysaccharides from *E.coli* (Sigma-Aldrich, #L2880) for 24 hr at 37⁰C and 5% CO_2_. For mouse M2 macrophages, RAW 264.7 cells were pretreated with 20 ng/ml recombinant mouse IL-4 (R&D Systems, #NBP2-76270) for 24 hr at 37⁰C and 5% CO_2_. For human M1/M2 macrophages, THP-1 cells were pretreated with 25 ng/ml phorbol 12-myristate 13-acetate (PMA, Sigma-Aldrich, #P1585) for 24 h at 37⁰C with 5% CO_2_, and then treated with 20 ng/ml IFN-γ (PeproTech, #300-02) and 20 ng/ml IL-4 (PeproTech, #200-04) for an additional 24 h to induce M1 and M2 macrophage polarization, respectively. For senescent macrophages, RAW cells or PMA-treated THP-1 cells were pretreated with 250 nM doxorubicin (CST, #5927S) for 24 hr at 37⁰C with 5% CO_2_, and then doxorubicin was removed by three washes with complete medium. Washed cells were cultured in the complete medium for RAW 264.7 cells and THP-1 macrophages, respectively, as described above for another 10 days before treatment with DGLA. Cell killing assays were performed using naïve, M1/M2-polarized or senescent macrophages in the presence of DGLA (Cayman, #90230) at a final concentration of 100 µM or vehicle control (ethanol). After incubation with DGLA or ethanol for 72 h at 37⁰C and 5% CO_2_, macrophage viability was determined using cell counting kit-8 (CCK-8), as described above.

#### Blue-red-green cell killing assay

Cells were plated in a 24-well plate and analyzed using the Abcam Apoptosis/ Necrosis Assay Kit (blue, green, red) (ab176749) to detect apoptotic, necrotic, and healthy cells. Prior to staining, NS or SEN(IR) cells were treated with DGLA or vehicle. Kit components were prepared according to manufacturer protocols, with volumes adjusted for a 24-well plate. Briefly, the cells were washed 2x with assay buffer and then replaced with staining solution containing 7-AAD, Cytocalcein violet 450, and Apopxin Green and then incubated at 37°C for 30 minutes. Cells were then washed with assay buffer and imaged on a MICA Microhub (Leica).

#### Bodipy-C11

BODIPY-C11 staining was conducted by making a 1.5 mM stock of BODIPY 581/591 C11 reagent (ThermoFisher, #D3861) in 100% ethanol, and then diluting 1:1000 to 1.5 µM in phenol red-free media. Media on the cells was aspirated off and replaced with BODIPY-C11 media, followed by a 20 minute incubation at 37°C and then imaging on a MICA Microhub (Leica). For 7 day analyses, media was then replaced with BODIPY C11-free growth media for and imaged 7 days later. Green and red channel fluorescence intensities were background normalized and quantified using ImageJ and presented as ratios of oxidized (green) to reduced (red).

#### Iron measurement (FerroOrange)

Detection of iron levels by FerroOrange staining was conducted by creating a 1 mM concentration of FerroOrange reagent (Dojindo), diluting 1:1000 to 1 uM in clear media, incubating the plate at 37°C for 30 minutes, washing 2x with PBS, and then images were captured in PBS on a Mica Microhub. Background subtracted image intensities per cell were calculated using ImageJ.

#### Sytox positivity assay

SYTOX Green staining was conducted in a 96-well plate using a concentration of 5 µM in clear media and read using a BioTek Cytation 5 Cell Imaging Multimode Reader, followed by addition of Hoechst. Cell death was expressed as percentage of Sytox positive nuclei relative to Hoechst positive nuclei.

#### Western blots

Protein lysate samples were generated by plating IMR-90s in 6-well plates. Sen(IR) samples were generated by irradiating cells with 10 Gy and then allowing them to sit for at least seven days prior to collection. MiDAS samples were generated by treating IMR-90s with 500 nm of antimycin A in DMSO for at twelve days. Media was aspirated followed by washing twice with ice-cold PBS. Cells were then scraped in RIPA lysis buffer (ThermoFisher, #89900) and homogenized by sonication. Protein levels were measured by Pierce BCA protein assay (ThermoFisher #23227). Samples containing 30 µg protein were then loaded into Criterion TGX Precast 12+12 well comb 4-15% polyacrylamide gels (Bio-Rad, #5671083). Precision Plus Protein Kaleidoscope Ladder (Bio-Rad, #161-0375) was loaded alongside the samples. The gel was run in a Bio-Rad Criterion Cell in 1x Tris/Glycine/SDS Buffer (Bio-Rad, #1610732). After running the gel, proteins were transferred to a nitrocellulose membrane (Bio-Rad, #1620115) in a Bio-Rad Criterion Blotter at 4°C at 30V overnight. Following transfer, protein transfer was visualized with Ponceau stain and subsequently blocked for 1hr in EveryBlot Blocking Buffer (Bio-Rad, #12010020). Membranes were then incubated at 4°C with primary antibodies diluted in EveryBlot Blocking Buffer (Bio-Rad) overnight. Antibodies used include the following: (1:1000, GPX4, Abcam, ab120566), (1:1000, ACSL4, ab155282, Abcam), (1:1000 LMNB1, Abcam, #ab16048), (1:1000, p16^INK4a^, Abcam, #ab108349). Following incubation with primary antibodies, nitrocellulose membranes were washed with PBST and incubated for 1 hr with secondary antibodies. Secondary ntibodies used: (1:20,000 Goat anti-rabbit IgG Alexa Flour Plus 488, Invitrogen, #A32731), (1:20,000 Goat anti-mouse IgG Alexa Flour Plus 800, Invitrogen, #A32730), (Goat anti-rabbit IgG Alexa Flour Plus 647, Invitrogen, #A32733). Membranes were then imaged using the Invitrogen iBright 1500 Imaging System. Following imaging of proteins of interest, actin was detected by incubating nitrocellulose membrane with primary antibody (1:5000, Actin, Abcam, #ab8226) for 30 minutes at room temperature, followed by washes in PBST and an hour in secondary antibody (1:20,000 Goat anti-mouse IgG Alexa Flour Plus 800, Invitrogen, A32730).

#### Immunofluorescence

IMR-90 fibroblasts were seeded in glass 8-well chamber slide (Thermo-Scientific, #154534) at 40k cells per well. Cells were allowed to grow and adhere for 72 h at 37C and 3% O2. Cells were irradiated at 10 Gy, followed by incubation at 37 C 3% O2 for 7 days. 5 days following irradiation, non-senescent IMR-90s were seeded at 40K/well in remaining wells of 8-well chamber slide to serve as controls. Cells were again allowed to adhere and grow for 72 hours at 37°C 3% CO2. 10 days following irradiation, half of non-senescent cells were treated with 1 µM RSL3 (Tocris, #6118) for 3 hours, then all cells were fixed in 10% phosphate buffered formalin (Azer Scientific, #PFNBF-240) for 10 min at room temperature. Cells were washed 3x with PBS for 5 min per wash. Cells were then permeabilized with 0.1% Triton-100X (Sigma, 9002-93-1) for 15 min at room temperature. Cells were washed 3x with PBS for 5 min per wash. Cells were then blocked with 10% normal goat serum blocking solution (Invitrogen, #50062Z) for 1 h at room temperature. Cells were then washed once with PBS for 5 min. Cells were incubated with primary antibody 4-HNE-adduct antibody (Invitrogen, #MA5-27570, 1:500) overnight at 4°C. Primary antibody was washed off with PBS 3x for 5 min per wash. Secondary antibody (AF488 goat anti-mouse, Invitrogen, #A11001, 1:1000) was added in a 0.1% BSA solution. This solution was added to cells for 1 h at room temperature covered from light. Secondary antibody solution was washed 3x with PBS for 5 min per wash, followed by aspiration and addition of mounting media with DAPI (Vector Laboratories, H-1200-10) and a glass coverslip placed on top. Slides were allowed to cure for 1 h according to manufacturer instructions and imaged using brightfield microscopy on a MICA at 10X magnification. Quantification of individual cell intensity normalized to background was performed in ImageJ.

#### Quantitative real-time PCR

RNA was isolated from animal tissue using a Zymogen Direct-zol RNA Minprep Kit (Zymo # R2072), mouse adipose tissue using the RNeasy Lipid Tissue Mini Kit (Qiagen) and from cultured cells using the Isolate II RNA Mini Kit (Meridian Biosciences). Isolated RNA reverse transcribed into cDNA using a High-Capacity cDNA Reverse Transcription Kit (ThermoFisher #4368813). Quantitative real time PCR was conducted using synthesized cDNA, (ThermoFisher #4352341E) or human actin (ThermoFisher #4326315E), and the corresponding custom or Taqman assay, listed in **Supplemental Table S1**.

#### Total RNA Isolation and Quantitative RT-PCR for macrophages

THP-1 cells were first pretreated with 25 ng/ml phorbol 12-myristate 13-acetate (PMA, Sigma-Aldrich, #P1585) for 24 h at 37⁰C with 5% CO_2_, and then treated with 20 ng/ml IFN-γ (PeproTech, #300-02) or 20 ng/ml IL-4 (PeproTech, #200-04) for another 24 hr to induce M1 and M2 macrophage polarization, respectively. Differentiated macrophages were then untreated or treated with vehicle ethanol or 100 µM DGLA (Cayman, #90230) for 24 h. Total cellular RNA was isolated using Monarch’s Total RNA Miniprep kit (New England Biolabs, #T20105) following manufacturer’s instructions. Quantitative RT-PCR was performed on the Roche LightCycler 480 real-time PCR system with Luna Universal Master Mix (Cat. No. M3003E; New England Biolabs) and primers for ARG-1, IL-10, CD38, LGALS3, TNF-α, and GAPDH.

### LIPID QUANTITATION ANALYTICAL METHODS

#### Cell membrane fatty acid extraction and quantitation

Non-senescent (NS) and SEN(IR) IMR-90 were treated simultaneously with either 200 µM DGLA or control (BSA) at ∼80% confluency and collected after 2 days (∼5 x 10^6^ cells) by scrapping and transfer to 1.5 mL vials using 200 μL of deionized water. Vials were stored at −80°C until further analysis.

To lyse cells, cell samples were quick thawed at 37°C and transferred into a 16×100ml glass test-tube. Cell membrane was isolated by washing with 2 mL of 4 °C 0.9% sodium chloride solution thrice and centrifuging for 10 mins at 1500 g at 4 °C. Membrane lipids were extracted using a modified Folch method^72^, after addition of an internal standard (heptadecanoic acid C17:0 at 1μg/μL). This was followed by saponification with methanolic sodium hydroxide (0.5N) and methylation with 14% boron-trifluoride-methanol^73^. The supernatant containing the fatty acid methyl esters (FAMEs) was dried down under nitrogen, resuspended in 100 µl of hexane, transferred into amber GC vials and stored at −20 °C until the time of analysis.

FAMEs were analyzed using a CLARUS 650 gas chromatograph (Perkin Elmer, Boston MA) equipped with a 100m x 0.25mm i.d (film thickness 0.25µm) capillary column (SP-2560, Supelco) as described previously^74^. Peaks of interest were identified by comparison with authentic fatty acid standards (Nu-Chek Prep, Inc. MN) and expressed both as absolute concentration (μg/mg weight) as well as molar percentage (mol %) proportions of total fatty acids relative to the internal standard.

#### Quantitation of lipids by flow injection analysis quadrupole time of flight mass spectrometry (FIA-QTOF-MS)

Lipids were isolated and quantitated according to a commercial kit (Biocrates MxP Quant 500) with the following conditions. 20 µL samples were injected into an Agilent 1290 liquid chromatograph using the chromatographic conditions prescribed by the kit’s instructions. The eluant from the HPLC was connected to an Agilent 6550 iFunnel quadrupole time-of-flight/mass spectrometry (QTOF) mass spectrometer with a Dual AJS electrospray ion source in positive ionization mode. Analyses were performed using the following ionization parameters: gas temperature (TEM) 200°C; drying gas, 13 psi; Vcap, 5500V; nebulizer, 60 psi; nozzle voltage 1000 volts; sheath gas temperature 250°C; sheath gas flow 11 L/min; and a reference mass of 149.02332 m/z. MS1 acquisition was operated in the positive ion scanning mode for a mass range of 75-3200 m/z. Because Biocrates supports triple quadrupole mass spectrometers but not QTOF instruments, we collected the data in high-resolution scan mode not in multiple reaction monitoring (MRM) mode as specified in the kit. As a result, we relied on the resolving power of the TOF not MS/MS for detection and identification of the compounds.

#### Data preprocessing

Because the Agilent QTOF data is not supported by Biocrates, we developed a data processing procedure using Agilent MassHunter Quantitative Analysis software version 10.1, Build 10.1.7330. Biocrates includes internal standards and calibrations samples in the kit, so we quantified these compounds using the Biocrates standards and calibrations curves. The Agilent Quantitative Analysis software did peak picking, integration, calibration curves and quantitative calculations automatically for most compounds, but some manual integration was required for low level metabolites. Because we used the Biocrates sample preparation kit with a QTOF there are some differences from the standard Biocrates data from a triple quadrupole mass spectrometer. The QTOF allowed us to collect quantitative targeted and untargeted metabolic profiles simultaneously. In FIA mode there is no separation using the Biocrates method because there is no column. Because we were not using MS/MS, the isomers of the individual DG’s and TG’s appear as one peak and could not be differentiated. The QTOF has high mass resolution, allowing identification of molecular formulas. Thus names of the DGs and TGs were changed to include the molecular formulas not the chain lengths as in the Biocrates software. For example TG(14:0_32:2) in the Biocrates database became TG(C_47_H_82_O_6_) in our database. Peak areas and standard-derived were normalized to total protein or cell number.

### ANIMAL MODEL ANALYSES

#### Whole body luminescence

22-24 month old p16-3MR mice^75^ were shaved, injected i.p. with RediJect Coelentrazine (PerkinElmer, # 760506) into the lower right of the peritoneal cavity, and luminescence was measured using an IVIS Lumina S5 Imaging System, and imaged again 2 weeks after DGLA-EE administration. All luminescence measurements were normalized to luminescence measurements from a phantom mouse.

#### Murine tissue function serum analyses

##### Free Fatty Acids (FFA) assay

Free Fatty Acids were measured in isolated mouse serum according to manufacturer instructions (Cayman, #700310). **Aspartate Aminotransferase (AST) assay.** The Aspartate Aminotransferase assay was performed on isolated mouse serum following manufacturer instructions (Cayman, #701640). **Albumin ELISA.** Albumin levels were measured in serum using a commercial kit (Abcam, #ab235628) according to manufacturer instructions. **Blood urea.** Urea was measured in mouse serum using a commercial kit (Abcam, #ab83362) according to manufacturer instructions.

#### Murine pathological and histological assessments

##### Liver pathology

OCT-embedded frozen tissue sections were stained with hematoxylin-eosin, Masson’s trichrome and picrosirius red staining. The slides were examined by investigator who was blinded to the treatment and only the sample numbers were provided. For each animal, whole section fields at a magnification of 10x, 40x and 100x were examined for relevant features:

1. Grading criteria for steatosis: Hepatic steatosis was graded at low power based on the percentage of the liver section involved by steatosis including both macro- and micro-vesicular fat accumulation: Grade 0 = < 5% (normal); Grade 1 = 5 to 25%; Grade 2 = 26 to 50%; Grade 3 = 51 to 75%; Grade 4 = 76% to 100%). Macrovesicular steatosis includes both large lipid droplets (occupying the cytoplasm, displacing the nucleus to the periphery) and small lipid droplets of variable size (occupying the cytoplasm with the nucleus maintaining its central location). Microvesicular steatosis is characterized by innumerable tiny, relatively uniform lipid vacuoles that result in a bubbly appearance of the hepatocytes.
2. Grading criteria for inflammatory foci: Hepatic inflammation was graded (from 0 to 3) according to the magnitude of hepatic infiltrates of inflammatory cells recruited immune cells or inflammatory infiltrate (usually patchy and composed of mostly lymphocytes with occasional plasma cells) and the number and size of inflammatory foci (clusters of the recruited immune cells) as follows: 0 = no inflammation (no hepatic infiltrates of inflammatory cells, no inflammatory foci); 1 = Mild hepatic infiltrates of inflammatory cells and some small inflammatory foci; 2 = Moderate hepatic infiltrates of inflammatory cells and several different sizes of inflammatory foci; 3 = Severe diffuse hepatic infiltrates of inflammatory cells and large sizes of inflammatory foci; and 4 = Massive hepatic infiltrates of inflammatory patchy and many large sizes of inflammatory foci.
3. Grading criteria for liver fibrosis: Liver fibrosis was scored according to the P.S. staining and M.T. staining, combined with H&E staining as follows: 0 = No staining detected; 1 = thickened perivenular collagen and a few thin collagen septa; 2 = 1 + thin septa with incomplete bridging between portal regions; 3 = 1 + 2 + thin septa and extensive bridging; and 4 = 1 + 2 + 3 + thickened septa with complete bridging of portal regions and a nodular appearance.
4. Calculation of fibrotic area: 3 PS-stained fields per animal liver were imaged, threshholded, and fibrotic area was quantified using ImageJ. Means of the 3 fields were presented as percentage fibrotic area per animal liver.

##### Adipose histology

Adipose tissue sections were stained with H&E and 3 sections were analyzed for adipocyte area and cell number per field using Adiposoft^76^. Crown-like structures were quantified for the same images.

## Acknowledgements

CW and CK thank Judith Campisi for the mentorship, support, and guidance that made this manuscript possible. This work was supported by USDA-ARS cooperative agreement 58-8050-9-004 (CW, AG, XDW, NM, TZ, GD), the Glenn Foundation for Medical Research and AFAR Grant for Junior Faculty (CW), pilot awards from the Tufts Healthy Aging Initiative and The Jackson Laboratory Nathan Shock Center (CW). BM was supported by the SENS Research Foundation. SJ was supported by NIDDK T32 DK124170 (PI: AG). RK was supported by AG038070 and AG079753.

## Author contributions

Conceptualization, CW, CK, SJ, BM; Investigation, CK, SJ, BM, NK, YC, MV, IS, RT, GD, CG, XT, AR, AE, TZ, KK, SD, CW. Resources, RK, TZ, MV, JP; Writing – CW, BM, NK, SJ, CK, GD; Review and Editing, CW, MV, JP; Supervision, CW, RK, AG, YC, NM; Formal analysis, XDW, NM, GD, AR; Funding acquisition, CW, AG, RK.

