## Supplemental Figures 1-4 for "Fatty acid desaturation guides cellular decisions between ferroptosis and cellular senescence"

**Supplemental Information**

**Supplemental Figure Legends**

**Figure S1. Altered PUFA desaturation in senescent cells, related to Figure 1.**

**A-C.** Cellular membranes were extracted from IR-induced senescent [SEN(IR)] or mock-irradiated non-senescent (NS) cells and analyze by HPLC for the indicated fatty acids.

A. Relative composition of saturated (SFA), monounsaturated (MUFA), or polyunsaturated (PUFA) in membranes of NS (red) vs. SEN(IR) (blue) cells.

B. Omega-6 fatty desaturation rates. (Left panel). Delta-6-desaturase (D6D) activity, as measured by DGLA:LA ratios, and (right panel) delta-5-desaturase (D5D) activity, as measured by AA:DGLA ratios, in NS and SEN(IR) cells.

C. Omega-3 fatty desaturation rates. (Left panel). Delta-6-desaturase (D6D) activity, as measured by SDA:ALA ratios, and (right panel) delta-5-desaturase (D5D) activity, as measured by EPA:SDA ratios, in NS and SEN(IR) cells.

D. Free fatty acid (FFA) desaturation rates in NS, SEN(IR), and MiDAS cells. (Left panel) Delta-6-desaturase (D6D) activity, as measured by DGLA:LA ratios, and (right panel) delta-5-desaturase (D5D) activity, as measured by AA:DGLA ratios, in NS and SEN(IR) cells.

**E-G**. Esterified fatty acids from NS or SEN(IR) IMR-90 fibroblasts were extracted 9 days after IR (or mock IR) and analyzed by mass spectrometry using a commercial kit (Biocrates).

E. D6D (left) and D5D (right) activity was estimated for phosphatidylcholines (PC) using 34:3 to 34:2 ratios for D6D and 34:4 to 34:3 for ratios for D5D.

F. D6D (left) and D5D (right) activity was estimated for lysophosphatidylcholines (LPC) using 20:3 to 18:2 ratios for D6D and 20:4 to 20:3 for ratios for D5D.

G. D6D (left) and D5D (right) activity was estimated for cholesterol esters (DE) using 20:3 to 18:2 ratios for D6D and 20:4 to 20:3 for ratios for D5D.

H. NS (PD ~32) and replicatively senescent SEN(REP) (PD ~70) IMR-90 fibroblasts were treated with 100 μM DGLA for 72 h and cell survival was measured by CCK-8 assay.

I.  Primary human hepatic stellate cells (HSteC) were induced to senesce by transduction with with a lentivirus containing the RasV12 oncogene [SEN(RAS)] or 500 nM antimycin A (MiDAS) for 7 days/. Empty vector cells treated with DMSO served as the non-senescent (NS) control. Cells were then treated with 100 μM DGLA for 72 h and cell survival was measured by CCK-8 assay normalized to groups treated with BSA.

J.  Mesenchymal stem cells (MSC) were induced to senesce by cultivation in 500 nM antimycin A (MiDAS) for 7 days, and cells treated with DMSO served as the non-senescent (NS) control. Cells were then treated with 100 μM DGLA for 72 h and cell survival was measured by CCK-8 assay normalized to groups treated with BSA.

K.  HepG2 cells were induced to senesce by culture in the presence of 10 μM Nutlin-3a or palbociclib (Palbo) for 7 days, followed by co-treatment with 100 μM DGLA for 72 h.  Cell survival was measured by CCK-8 assay normalized to groups treated with BSA.

L.  NS or SEN(IR) IMR-90 fibroblasts were treated with the indicated doses of gamma-linolenic acid (GLA) for 72 h.  Cell survival was measured by CCK-8 assay normalized to groups treated with BSA.

M.  NS or SEN(IR) IMR-90 fibroblasts were treated with the indicated doses of dihomo-alpha-linolenic acid (DALA) for 72 h.  Cell survival was measured by CCK-8 assay normalized to groups treated with BSA.

N.  NS or SEN(IR) IMR-90 fibroblasts were treated with the indicated doses of Mead's acid for 72 h.  Cell survival was measured by CCK-8 assay normalized to groups treated with BSA.

O.  NS or SEN(IR) IMR-90 fibroblasts were treated with the indicated doses of the Steroyl-CoA desaturase (SCD) inhibitors CAY10566 (left) or A-939572 (right) for 6 days.  Cell survival was measured by CCK-8 assay normalized to groups treated with DMSO.

P. NS or SEN(IR) IMR-90 fibroblasts were treated with the indicated doses of the fatty acid synthase (FASN) inhibitors CMS121 (left) or TVB-3664 (right) for 6 days. Cell survival was measured by CCK-8 assay normalized to groups treated with DMSO.

Q. NS or SEN(IR) IMR-90 fibroblasts were treated with the indicated doses of the delta-6 desaturase inhibitor sc-26196 for 6 days. Cell survival was measured by CCK-8 assay normalized to groups treated with DMSO.

**Figure S2. DGLA does not kill senescent cells by apoptosis, related to Figure 2.**

A. NS or SEN(IR) IMR-90 fibroblasts were treated with the indicated doses of 8-hydroxy-octanoic acid (8-HOA) for 72 h. Cell survival was measured by CCK-8 assay normalized to groups treated with DMSO.

B. NS and SEN(IR) cells were treated with 100 μM DGLA for 48 h, followed by membrane isolation, extraction of fatty acids, and measurement by HPLC.

C. Esterified fatty acids from NS or SEN(IR) IMR-90 fibroblasts were extracted 9 days after IR (or mock IR) treated with BSA or 100 μM DGLA for 48 h and analyzed by mass spectrometry using a commercial kit (Biocrates). All detected lipid species containing 3 or more double bonds are presented in the heat map. * = $p<0.05$, two-way ANOVA.

**Figure S3. Ferroptosis inducers selectively kill senescent cells, related to Figure 3.**

A. NS or SEN(IR) primary human hepatic stellate cells or mesenchymal stem cells (MSCs) were analyzed for labile iron content by FerroOrange staining.

B. NS or SEN(IR) IMR-90 fibroblasts were treated with the indicated doses of ML-210 for 18 h and analyzed by CCK-8 assay.

C. NS or SEN(DOXO) IMR-90 fibroblasts were treated with the indicated doses of RSL3 for 18 h and analyzed by CCK-8 assay.

D. NS or SEN(DOXO) IMR-90 fibroblasts were treated with the indicated doses of erastin for 18 h and analyzed by CCK-8 assay.

E. NS or SEN(IR) murine astrocytes were treated with the indicated doses of ML-210 for 18 h and analyzed by CCK-8 assay.

F. NS or SEN(IR) murine astrocytes were treated with the indicated doses of RSL3 for 18 h and analyzed by CCK-8 assay.

G. NS or SEN(IR) HepG2 cells were treated with the indicated doses of ML-210 for 18 h and analyzed by CCK-8 assay.

H. NS or SEN(IR) HepG2 cells were treated with the indicated doses of RSL3 for 18 h and analyzed by CCK-8 assay.

I. NS or SEN(IR) IMR-90 fibroblasts were treated with 1 μM RSL3 and percent Sytox positive cells (normalized to end point Hoechst) were calculated for 3 wells at the indicated timepoints.

J. NS or replicatively senescent [SEN(REP)] IMR-90 fibroblasts were treated with 1 μM RSL3 and percent Sytox positive cells (normalized to end point Hoechst) were calculated for 3 wells at the indicated timepoints.

K. NS or SEN(IR) IMR-90 fibroblasts were treated were treated with 3 μM FIN-56 and percent Sytox positive cells (normalized to end point Hoechst) were calculated for 3 wells at the indicated timepoints.

**Figure S4. DGLA-induced loss of senescence markers is not due to macrophages, related to Figure 4.**

A. NS or SEN(IR) IMR-90 fibroblasts were treated with 0, 100 μM, or 200 μM DGLA ethyl ester (DGLA-EE) and cell survival was measured 72 h later by CCK-8 assay.

B. Adipocyte numbers per field in visceral white adipose tissue were calculated for 3 fields per animal for aged male and female mice treated with DGLA-EE or vehicle.

C.  Average percent steatotic area for aged male and female aged mice treated with DGLA-EE or vehicle.

D.  RNA from liver tissue from male and female mice treated with DGLA-EE or vehicle was analyzed by qPCR for *Lgals3*.

E.  RNA from liver tissue from male and female mice treated with DGLA-EE or vehicle was analyzed by qPCR for *Cd68*.

F.  RAW264.7 murine macrophages were induced to undergo M1 polarization with 20 ng/ml recombinant mouse IFN-γ and 100 ng/ml LPS, M2 polarization with 20 ng/ml recombinant mouse IL-4, or induced to senesce by incubation with 250 nM doxorubicin for 24 h and allowed to recover for 10 days.  Untreated RAW264.7 served as controls.  Cells were then treated with either vehicle (EtOH) or 100 μM DGLA or 72 h, and viability was determined by CCK-8 assay.

G.  Control (CTL), M1, or M2 Thp-1 macrophages were treated with DGLA for 72 h, RNA was extracted, and gene expression of *LGALS3* was analyzed by qPCR, normalized to actin.

H.  Control (CTL), M1, or M2 Thp-1 macrophages were treated with DGLA for 72 h, RNA was extracted, and gene expression of *ARG1* was analyzed by qPCR, normalized to actin.

I.  Control (CTL), M1, or M2 Thp-1 macrophages were treated with DGLA for 72 h, RNA was extracted, and gene expression of *IL10* was analyzed by qPCR, normalized to actin.

J.  Control (CTL), M1, or M2 Thp-1 macrophages were treated with DGLA for 72 h, RNA was extracted, and gene expression of *TNF* was analyzed by qPCR, normalized to actin.

K.  Control (CTL), M1, or M2 Thp-1 macrophages were treated with DGLA for 72 h, RNA was extracted, and gene expression of *CD38* was analyzed by qPCR, normalized to actin.

L.  RNA from adipose tissue from male and female mice treated with DGLA-EE or vehicle was analyzed by qPCR for *Cd38*.

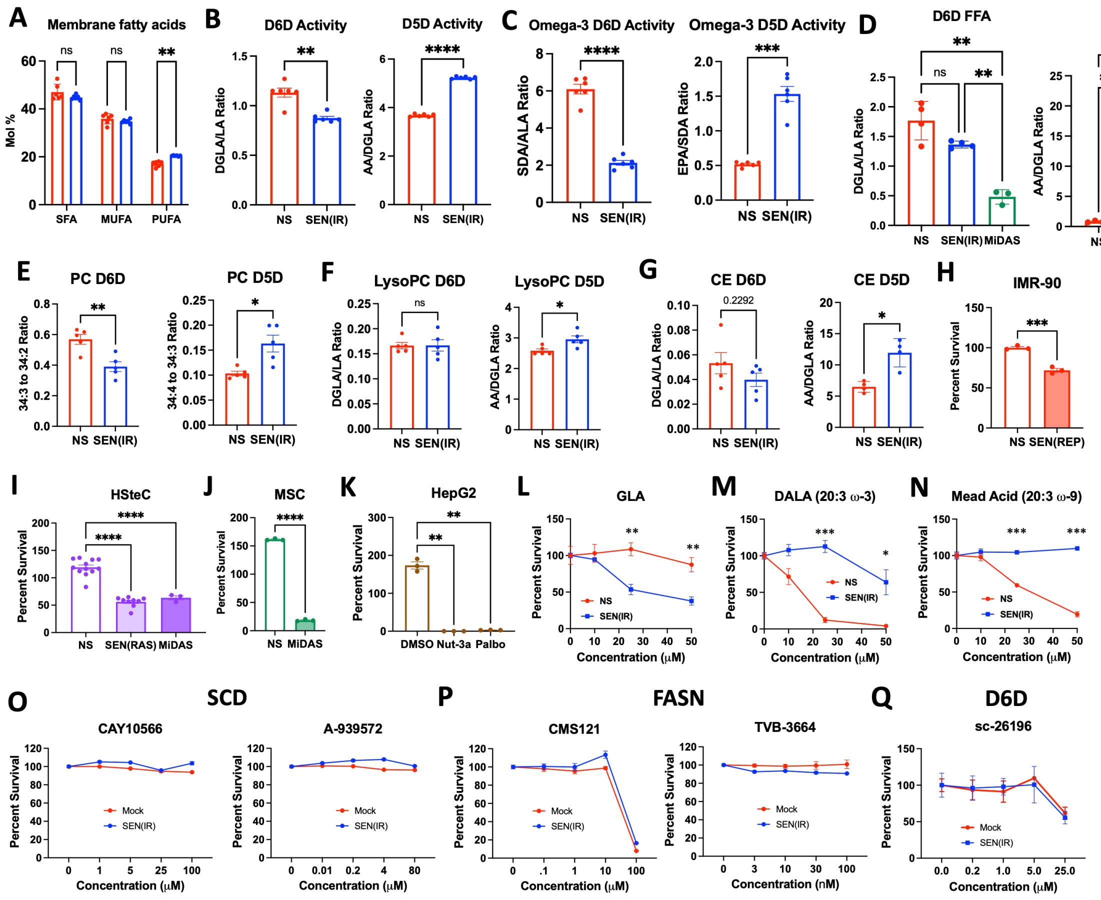
A Membrane fatty acids
B D6D Activity
D5D Activity
C Omega-3 D6D Activity
Omega-3 D5D Activity
D D6D FFA
D5D FFA
E PC D6D
PC D5D
F LysoPC D6D
LysoPC D5D
G CE D6D
CE D5D
H IMR-90
I HSteC
J MSC
K HepG2
L GLA
M DALA (20:3 ω-3)
N Mead Acid (20:3 ω-9)
O SCD
CAY10566
A-939572
P FASN
CMS121
TVB-3664
Q D6D
sc-26196

Figure S1.

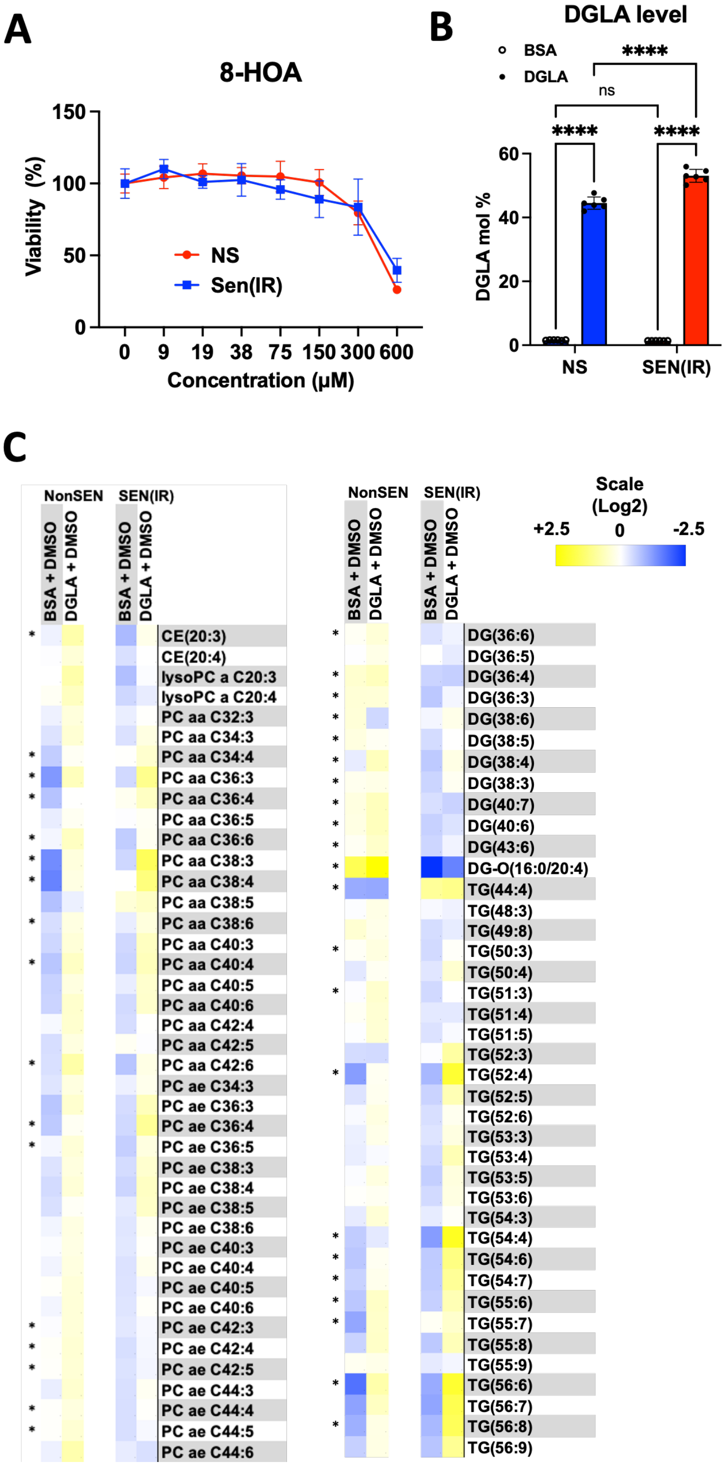
A
8-HOA
Viability (%)
150
100
50
0
NS
Sen(IR)
0 9 19 38 75 150 300 600
Concentration (μM)
B
DGLA level
BSA
DGLA
****
ns
****
****
DGLA mol %
60
40
20
0
NS
SEN(IR)
C
NonSEN
SEN(IR)
BSA + DMSO
DGLA + DMSO
CE(20:3)
CE(20:4)
lysoPC a C20:3
lysoPC a C20:4
PC aa C32:3
PC aa C34:3
PC aa C34:4
PC aa C36:3
PC aa C36:4
PC aa C36:5
PC aa C36:6
PC aa C38:3
PC aa C38:4
PC aa C38:5
PC aa C38:6
PC aa C40:3
PC aa C40:4
PC aa C40:5
PC aa C40:6
PC aa C42:4
PC aa C42:5
PC aa C42:6
PC ae C34:3
PC ae C36:3
PC ae C36:4
PC ae C36:5
PC ae C38:3
PC ae C38:4
PC ae C38:5
PC ae C38:6
PC ae C40:3
PC ae C40:4
PC ae C40:5
PC ae C40:6
PC ae C42:3
PC ae C42:4
PC ae C42:5
PC ae C44:3
PC ae C44:4
PC ae C44:5
PC ae C44:6
Scale
(Log2)
+2.5
0
-2.5
DG(36:6)
DG(36:5)
DG(36:4)
DG(36:3)
DG(38:6)
DG(38:5)
DG(38:4)
DG(38:3)
DG(40:7)
DG(40:6)
DG(43:6)
DG-O(16:0/20:4)
TG(44:4)
TG(48:3)
TG(49:8)
TG(50:3)
TG(50:4)
TG(51:3)
TG(51:4)
TG(51:5)
TG(52:3)
TG(52:4)
TG(52:5)
TG(52:6)
TG(53:3)
TG(53:4)
TG(53:5)
TG(53:6)
TG(54:3)
TG(54:4)
TG(54:6)
TG(54:7)
TG(55:6)
TG(55:7)
TG(55:8)
TG(55:9)
TG(56:6)
TG(56:7)
TG(56:8)
TG(56:9)

**Figure S2.**

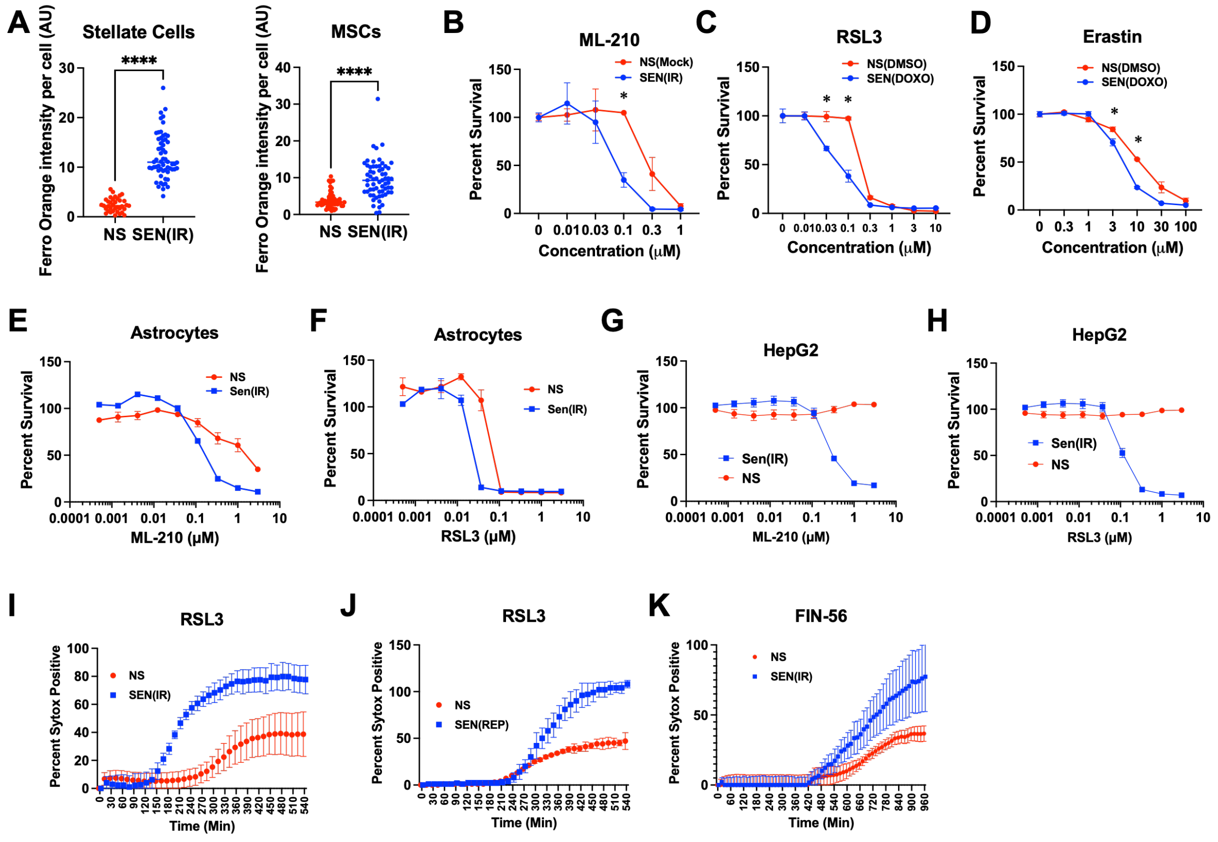

A
Stellate Cells
MSCs
Ferro Orange intensity per cell (AU)
****
NS
SEN(IR)
B
ML-210
NS(Mock)
SEN(IR)
Percent Survival
Concentration (μM)
C
RSL3
NS(DMSO)
SEN(DOXO)
D
Erastin
E
Astrocytes
NS
Sen(IR)
ML-210 (μM)
F
RSL3 (μM)
G
HepG2
H
I
Percent Sytox Positive
Time (Min)
J
SEN(REP)
K
FIN-56


**Figure S3.**

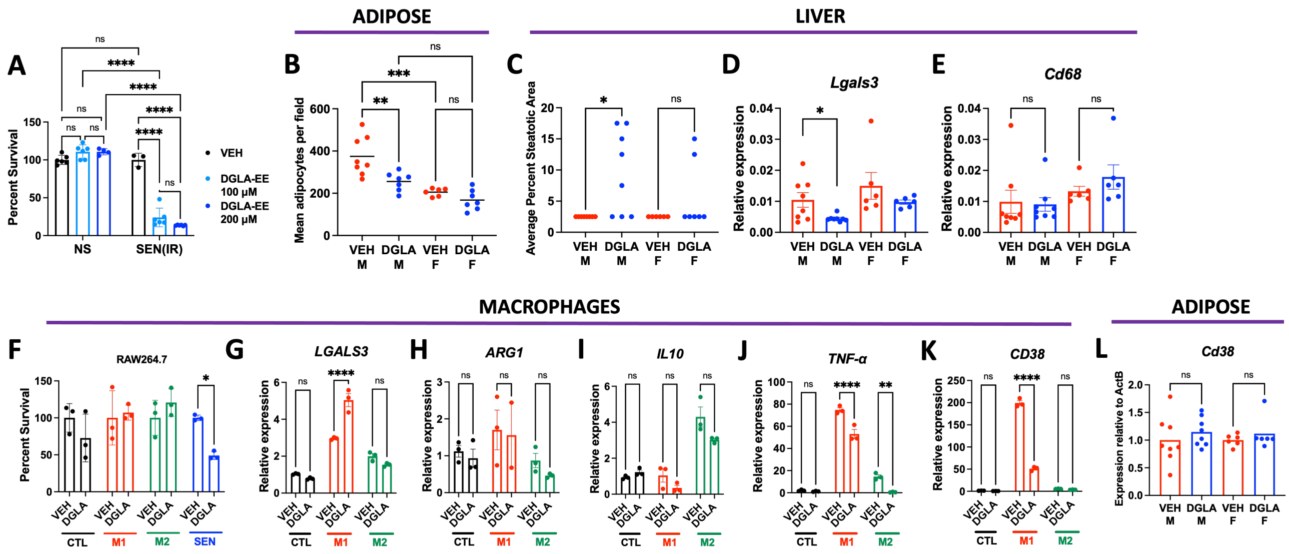
ADIPOSE
LIVER
A
B
C
D
E
Lgals3
Cd68
Percent Survival
NS
SEN(IR)
VEH
DGLA-EE 100 μM
DGLA-EE 200 μM
Mean adipocytes per field
Average Percent Steatotic Area
Relative expression
MACROPHAGES
ADIPOSE
F
G
H
I
J
K
L
RAW264.7
LGALS3
ARG1
IL10
TNF-α
CD38
Cd38
Expression relative to ActB
CTL
M1
M2
SEN

Figure S4.
